# Multifaceted mRNA analysis using programmed RNA cleavage by *Mucilaginibacter paludis* Argonaute

**DOI:** 10.1101/2025.07.01.662632

**Authors:** Lizhi Liu, Eric A. Hunt, Eric J. Wolf, Ivan R. Corrêa, Erbay Yigit, Sebastian Grünberg

## Abstract

Argonaute (Ago) endonucleases are ubiquitous across all domains of life and, like Cas nucleases, can be programmed by guide oligonucleotides to target and cleave specific nucleic acid sequences. However, unlike Cas enzymes, Agos are not restricted by sequence requirements such as protospacer adjacent motifs (PAMs) enhancing their utility as molecular tools. This study demonstrates diverse applications of the DNA-guided RNA-endonuclease *Mucilaginibacter paludis* Argonaute (Mpa Ago) for *in vitro* mRNA processing, e.g., 3’ end polishing, and for mRNA analysis in combination with liquid chromatography tandem mass spectrometry (LC-MS/MS). Applications of the latter include the comprehensive assessment of 5’ capping efficiency, 3’ poly(A) tail length and composition, one-pot multiplexed cleavage for RNA sequencing, modification detection, and fingerprinting.

## Main

Like CRISPR-Cas nucleases, Agos are conserved endonucleases that utilize nucleic acid guides to recognize, bind, and cleave their substrates. However, unlike Cas nucleases, Ago proteins are present across all domains of life^1–3^. In eukaryotes, Agos are loaded with small viral RNAs, endogenous miRNAs, or piRNAs, functioning in RNA interference (RNAi) pathways as components of the RNA-induced silencing complex (RISC)^4–8^. They play crucial roles in antiviral defense, posttranscriptional regulation of gene expression, and suppression of transposon activity^9–11^.

Prokaryotic Agos (pAgos) are classified into long and short based on their domain architecture^2, 3^. Long pAgos consist of six domains (N-terminal, L1, PAZ, L2, MID, and PIWI) and share structural similarities with eukaryotic Agos (eAgos). However, some long pAgos are catalytically inactive due to disruptions in their catalytic tetrad, DEDX (X = D/H/N/K), which coordinate divalent metal cations, often Mg^2+^, that are crucial for activity^1, 2, 12–14^. Short pAgos contain only the MID and PIWI domains and lack the ability to cut nucleic acids due to mutations in the catalytic tetrad^15^. Short pAgos and catalytically inactive long pAgos often associate with other effector proteins which function in bacterial defense systems^16^. In contrast, active long pAgos have been reported to be directly involved in cellular defense against foreign genetic elements as well as in genome replication and recombination^12, 17–21^.

Active pAgos load short guides to form a nucleoprotein complex that subsequently binds and cleaves complementary target sequences between guide nucleotides 10 and 11, resulting in cleavage products with 3’-OH and 5’-phosphate ends^22^. Unlike eAgos, which exclusively bind RNA guides to cleave RNA targets, most known active long pAgos utilize short DNA guides to target DNA sequences^17, 18, 23–30^. Few pAgos have been reported to use RNA guides to cleave DNA^14, 31–34^, and still fewer using RNA guides to target RNA^33, 35, 36^. Recently, several pAgos have been reported to use DNA guides to preferentially cleave RNA over DNA^28, 37, 38^.

The ability to direct endonuclease activity to a specific location with short nucleic acid guides makes pAgos highly versatile and valuable molecular tools with many applications in biotechnology. When compared to Cas nucleases, the lack of sequence restrictions such as PAMs and the predominant use of short DNA guides versus RNA guides make pAgos a versatile and economical alternative, especially for in vitro applications. DNA-targeting pAgos are already used in the diagnosis of pathogens and disease related rare mutations, DNA assembly, super-resolution microscopy, and the detection of single and double stranded DNA sequences^39^. However, applications that utilize pAgos to specifically target RNA have not yet been reported.

In this work, we describe RNA-specific applications of MpaAgo, a DNA-guided pAgo that preferentially targets RNA substrates^37^. We demonstrate that MpaAgo can be used to efficiently cleave a target RNA at 24 sites simultaneously in a multiplex one-pot reaction, allowing for subsequent RNA sequencing using intact mass spectrometry with up to >90% sequence coverage. Custom guide mixtures generating characteristic RNA fragments enable RNA fingerprinting – a fast and unambiguous way to detect and identify RNAs by liquid chromatography. We also show that MpaAgo can be used in 5’ cap analysis of mRNAs, allowing for the precise determination and relative quantification of the cap incorporation in a workflow that can be adapted to virtually any 5’ untranslated region (UTR) sequence and can be performed in under one hour. Lastly, we demonstrate that MpaAgo can be guided to and specifically cleave a sequence embedded in the poly(A) tail of an *in vitro* transcribed mRNA to generate mRNA populations with homogenized 3’ ends, removing non-templated additions introduced by RNA polymerases. This also allows for the accurate analysis of the poly(A) tail length and identification of unwanted potential extensions.

## Results

### MpaAgo efficiently and specifically cleaves mRNA

To assess the potential of pAgos to manipulate therapeutically relevant RNA substrates, we sought to determine the specificity and cleavage efficiency of mRNA substrates by MpaAgo, a DNA-guided endoribonuclease from *Mucilaginibacter paludis*. MpaAgo has previously been described as a programmable endoribonuclease with strong guide-mediated substrate specificity that is capable of cleaving RNA substrates of up to 352 nt in length^37^. We tested site-specific RNA cleavage by MpaAgo on two different *in vitro* transcribed (IVT) mRNAs: pSG90 IVT RNA, which represents a shortened version of the replication-competent Sindbis virus genome^40^ (1678 nt), and Fluc (*Firefly luciferase*) IVT RNA (1766 nt). We designed 16 nt long 5’-phosphorylated guides targeting unique sequences in the pSG90 and Fluc IVT RNAs, spaced 100 or 72 nt apart throughout the complete mRNA sequences. When cleaving the pSG90 RNA with MpaAgo using single guides in separate reactions, we observed cleavage efficiencies between 71% and 92% compared to a control reaction containing only MpaAgo without guide (Fig. 1A). Compared to pSG90 RNA, MpaAgo displayed a higher cleavage efficiency of Fluc RNA, resulting in near complete cleavage of the substrate mRNA for most guides (Fig. 1B).

**Figure 1.**
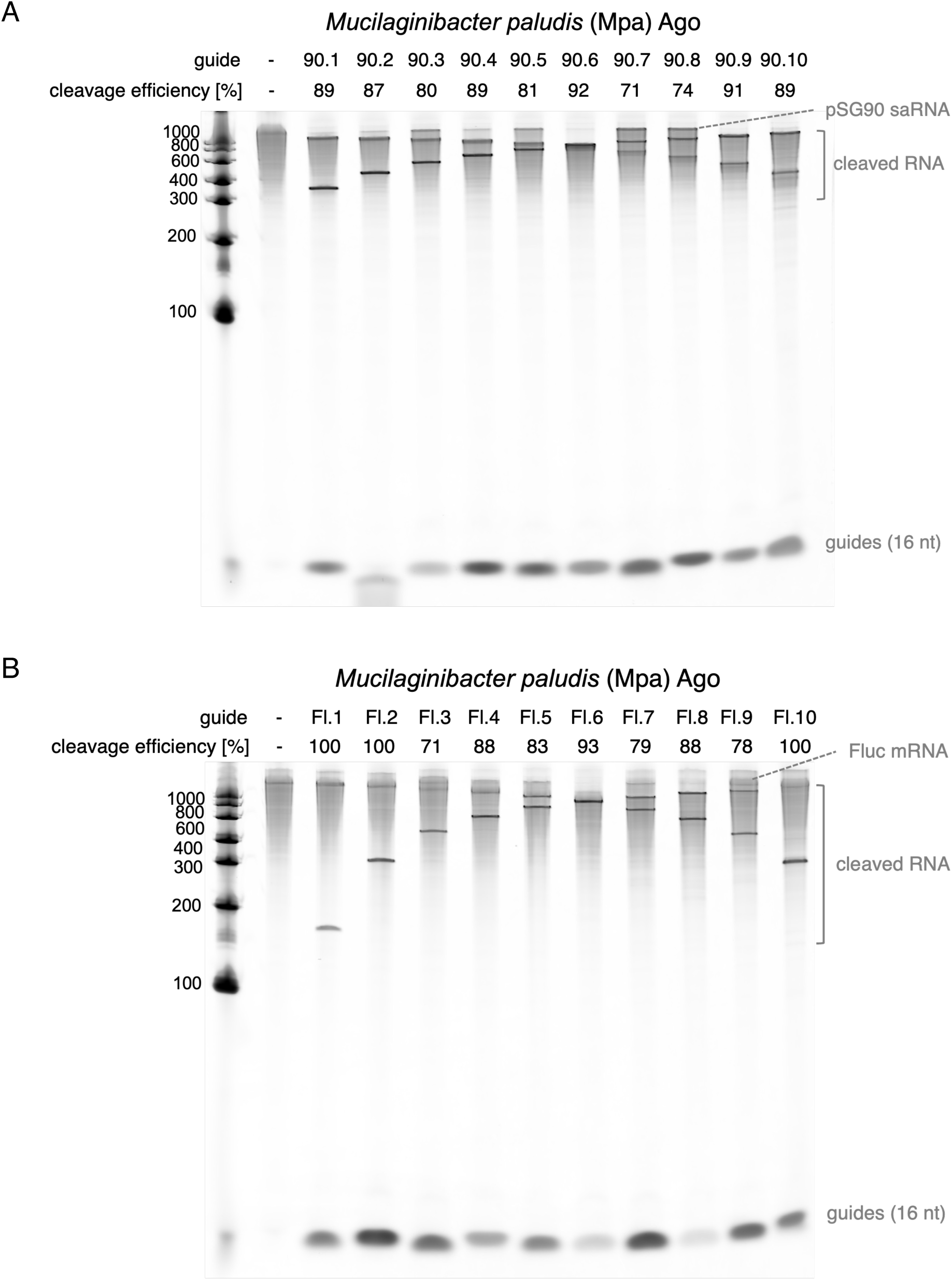
MpaAgo efficiently and specifically cleaves mRNAs. **(A)** pSG90 IVT RNA (∼1.7 kb) was subject to MpaAgo cleavage with single complementary DNA guides in ThermoPol® reaction buffer at 50°C for 15 minutes. The substrate:Ago molar ratio was 1:5, and the Ago:guide molar ratio was 1:5. 16 nt guides were 5’ phosphorylated and were designed for MpaAgo to cleave 100 nt apart from each other. Cleavage products were analyzed on a denaturing 6% TBE-urea gel stained with SYBR Gold. Guide identities and cleavage efficiencies (%) are indicated above the corresponding lanes. **(B)** Fluc IVT RNA (∼1.8 kb) was subject to MpaAgo cleavage with single complementary DNA guides. Cleavage reactions were performed under the same conditions as in (A). 16 nt guides for Fluc RNA were also 5’ phosphorylated but were designed for MpaAgo to cleave 72 nt apart from each other. Cleavage products were analyzed on a denaturing 6% TBE-urea gel stained with SYBR Gold. Guide identities and cleavage efficiencies (%) were indicated above the corresponding lanes.

To challenge MpaAgo further, we tested cleavage of a uridine-depleted version of the human erythropoietin (Epo) IVT RNA with a GC content (excluding the poly(A) tail) of 59% (pSG95; Fig. S1). The ability to cleave GC-rich substrates is a desirable property, as uridine depletion and/or modification has been shown to reduce immunogenicity and is commonly applied to RNA therapeutics design^41^. All but one guide enabled cleavage with efficiencies close to those observed in the other two tested mRNAs (between 71% to 89%); the outlier, guide 95.2, only cleaved ∼ 50% of the substrate, possibly due to local secondary structures that were not fully resolved under the conditions tested.

To assess the possibility of off-target cleavage by MpaAgo, we selected guides targeting sequences in Fluc RNA and used them to target pSG90 RNA. We did not detect cleavage, whether in the presence of guides (Fig. S2), further corroborating the highly specific nature of cleavage. Previous work reported that guide length may affect cleavage efficiency on short RNA substrates^37^. We repeated cleavage of Fluc IVT RNA with MpaAgo using 18 nt DNA guides and found similar cleavage efficiencies compared to that using 16 nt guides, with results varying between 61% and complete cleavage (Fig. S3).

To expand on the initial characterization of MpaAgo by Li et al.^37^, we tested the efficiency of MpaAgo to cleave long RNAs (> 1000 nt) at various temperatures and incubation times (Fig. S4A). We observed very little cleavage of the 1678 nt pSG90 IVT RNA at 4°C even after a 16-hour incubation (Fig. S4B). Increasing the temperature to 24°C and 37°C significantly increased cleavage efficiency, with much of the substrate being cut after 30 min incubation at 37°C. Cleavage by MpaAgo was highly effective at 50°C, with most of the substrate completely cleaved after 10 min. Importantly, both MpaAgo and a guide were required for specific cleavage as neither MpaAgo nor the guide alone induced cleavage of the substrate RNA (Fig. S4B). MpaAgo-independent degradation of the RNA substrate was observed at 37 or 50°C for 60 min or longer incubation times (Figs. S4A, right panel and S4B), likely due to Mg^2+^ catalyzed hydrolysis of phosphodiester bonds^42^.

We next tested if the DNA guides could be efficiently removed using DNA exonucleases, which would allow DNA-free downstream processes. *E. coli* exonuclease I, which is a 3’-5’ exonuclease with strong preference for single stranded DNA, readily degraded most of the DNA guide from the pAgo reaction (Fig. S5, lane 2). We hypothesize that the small amount of remaining guide may be bound to and thus protected by the MpaAgo protein.

### Multiplexed cleavage of mRNA

After confirming that pAgos specifically and efficiently cleave RNAs longer than 1000 nt, we proceeded to investigate whether MpaAgo could generate the corresponding cleavage fragments in a single reaction when provided with a pool of guides targeting multiple locations along an RNA (Fig. 2A). Consistent with the single cleavage reactions shown in Fig. 1A, incubating MpaAgo with two guides (90.1 and 90.2) resulted in specific cleavage of the full-length pSG90 RNA at two individual sites (Fig. 2B). The length of the resulting ∼100 nt cleavage fragment (Fig. 2B, lane 4) is consistent with the distance between the target sequences of the two guides. To completely cleave the full-length Fluc RNA into unique 72 nt fragments (plus a 29 nt 5’ fragment and an 81 nt 3’ fragment), we combined 24 16 nt DNA guides in a single reaction. Using a ratio of substrate to enzyme to total guide pool of 0.1:1:1, complete cleavage of Fluc IVT RNA was achieved (Fig. 2C, experiment 2). However, several fragments of sizes >72 nt were still visible, suggesting that not all guides enabled complete cleavage of their respective target sequences. The addition of 0.5% bovine serum albumin (BSA) to the reaction enhanced cleavage efficiency, collapsing the banding pattern to only three bands besides the unique 5’ and 3’ end fragments: the desired 72 nt fragments, one prominent band running at 144 nt, and one faint fragment with a size of ∼ 220 nt (Fig. 2C, experiment 4). The 144 nt and 220 nt bands likely represent fragments with one or two missing cleavages, indicating that cleavage was still incomplete. We then tested additional supplements to further increase cleavage efficiency. In the absence of BSA, the addition of an 1x enhancer mix containing 0.5 M betaine, 20 mM potassium glutamate, 20% glycerol, and 5% PEG 3350 to the reactions resulted in a similar fragment distribution as did the addition of 0.5% BSA (Fig. 2C, experiment 5). Combining the enhancer mix with 0.5% BSA resulted in the complete cleavage of Fluc mRNA into the programmed RNA fragment sizes (Fig. 2C, experiment 3). Cleaved fragments from the reactions containing the guide pool (experiments 2-5) were submitted to LC-MS/MS analysis, indicating sequence coverages between 79 and 88% (Fig. 2D and Fig S6). Interestingly, even the reaction without additives that showed only partial cleavage resulted in 79.2% Fluc mRNA sequence coverage (Fig. 2D and Fig. S6, experiment 2), suggesting that complete digestion of the substrate mRNA may not be a prerequisite for high mapping coverage.

**Figure 2.**
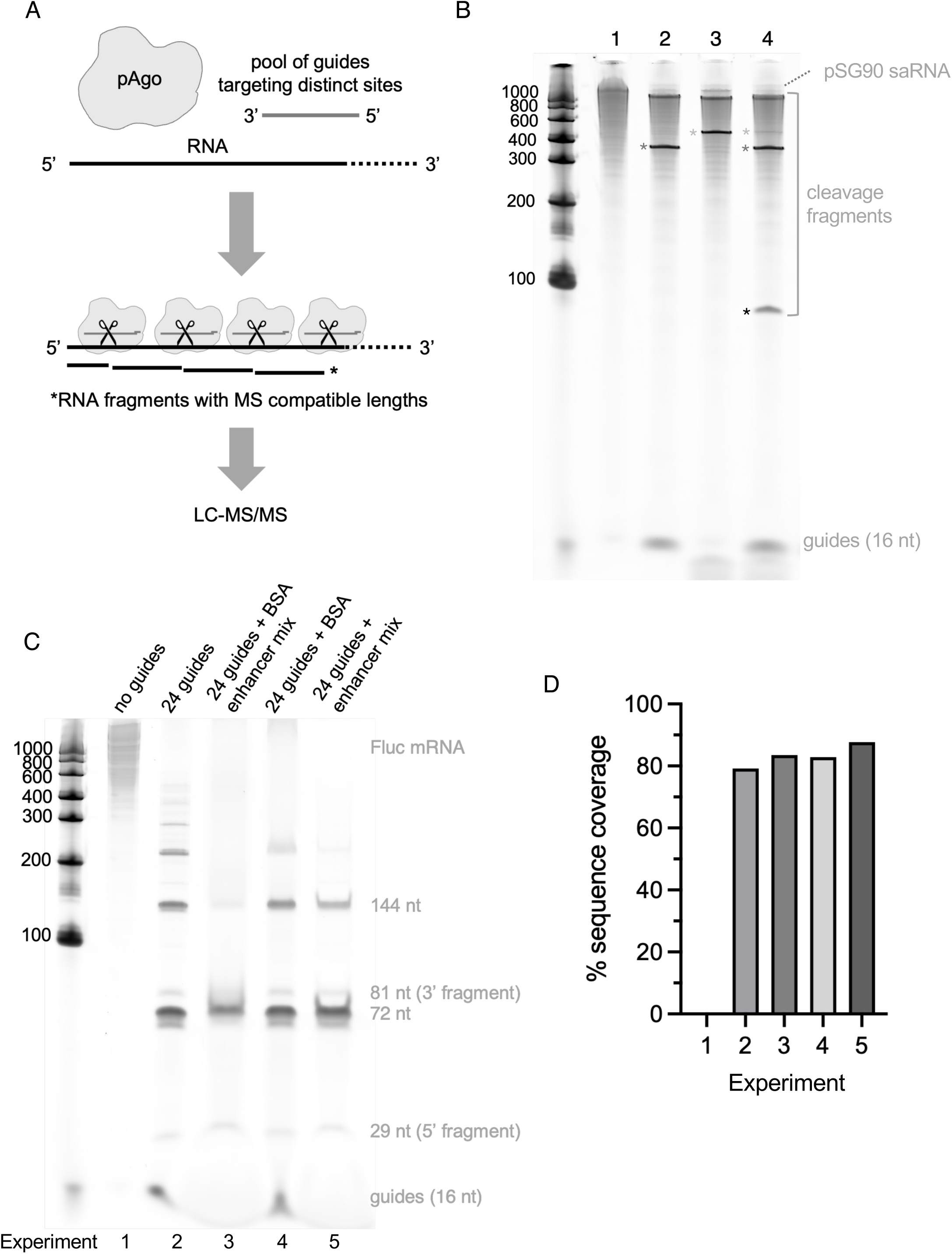
Guide multiplexing with MpaAgo produces RNA fragments suitable for LC-MS/MS analysis. **(A)** Schematic showing guide multiplexing with pAgo. Guides targeting different regions of an RNA are pooled and used in a single reaction to cleave an RNA into smaller fragments compatible with LC-MS/MS analysis. **(B)** pSG90 IVT RNA (∼1.7 kb) was subject to MpaAgo cleavage with single guides (lanes 2 and 3) or two 5’ phosphorylated guides in a single reaction (lane 4), at 50°C for 30 minutes. The substrate:Ago molar ratio was 1:5, and the Ago:guide molar ratio was 1:5. Cleavage products were analyzed on a denaturing 6% TBE-urea gel stained with SYBR Gold. The resulting short cleavage fragments are indicated by asterisks. The target sites for the two guides were 100 nt apart, producing a new 100 nt fragment (black asterisk) when used together. This fragment migrated faster than the 100 nt marker likely due to different buffer compositions or unresolved secondary structure elements. **(C)** Fluc IVT RNA (∼1.8 kb) was subjected to MpaAgo cleavage with a pool of 24 5’ phosphorylated guides, with or without 0.5% BSA or enhancer mix at 50°C for 30 minutes as indicated above the gel. The substrate:Ago molar ratio was 1:10, and the Ago:guide molar ratio was 1:1. Cleavage products were analyzed on a denaturing 6% TBE-urea gel stained with SYBR Gold. The distance between target sites of adjacent guides was 72 nt, such that complete cleavage would produce fragments of a uniform 72 nt length in addition to unique 5’ and 3’ fragments of 29 nt and 81 nt, respectively. Bands ≥ 72 nt represent incomplete cleavage fragments. **(D)** Cleavage products from (C) were analyzed by LC-MS/MS, and the sequence coverage of Fluc RNA (%) are shown in a bar graph. Experimental conditions: 1. no guides; 2. a pool of 24 guides, no additives; 3. 24 guides together with 0.5% BSA and the enhancer mix; 4. 24 guides with 0.5% BSA; and 5. 24 guides with the enhancer mix.

Recent data suggest that MpaAgo can also use longer DNA guides to cleave RNA^37^. We thus decided to use 18 nt guides to test if longer guides could increase the specificity of MpaAgo targeting and cleavage. While the 16 nt guides were selected to generate uniform 72 nt RNA fragments, the 18 nt guides were designed to generate laddering RNA fragments ranging from 69 to 81 nt and thus simplify oligonucleotide analysis by LC-MS/MS. Under conditions similar to those used with the 16 nt guides, Fluc and pSG90 IVT RNAs were incubated with MpaAgo and their respective 18 nt guide pools (Fig. S7A). Consistently, the addition of the enhancer mix significantly increased RNA site-specific cleavage, collapsing most of the RNA fragment sizes to around 60-80 nt (Fig. S7A, compare experiments 2 and 3, and experiments 5 and 6). Sequence coverage for pSG90 and Fluc mRNAs ranged from 78 to 95%, based on LC-MS/MS analysis (Figs. S7B and S7C).

To assess whether an RNA modification commonly used in therapeutic RNAs affect MpaAgo cleavage accuracy and efficiency, we incubated it with a Fluc IVT RNA in which all uridines were replaced with *N*1-methylpseudouridine (m^1^Ψ) and the same 18 nt guide pool as above. Gel electrophoresis revealed partially cleaved higher molecular weight fragments even in the presence of the enhancer mix (Fig. S8A), suggesting that m^1^Ψ may affect the cleavage efficiency. Even though cleavage of the fully m^1^Ψ-modified Fluc was not complete, the ensemble of identified cleavage products was sufficient to generate RNA mapping coverages of around 77% (without enhancer mix) and 86% (with enhancer mix), as determined by LC-MS/MS analysis (Figs. S8B and S8C), which were only marginally lower than those of unmodified Fluc (78 and 95%, respectively).

Overall, these data suggest that MpaAgo can be combined with multiple guides in a single reaction to cleave long RNA substrates into fragments of defined sizes, effectively reducing the complexity of the fragment pool (compared to non-specific RNA chemical cleavage or commonly used RNases with short recognition motifs) and enabling high sequence mapping coverage using oligonucleotide LC-MS/MS analysis.

We next investigated whether pooled guide DNAs could be employed to simultaneously target multiple RNA species in a one-pot reaction. The resulting cleavage products then serve as species-specific molecular fingerprints, which can be detected by LC-MS/MS to include or exclude RNA species. In reactions containing Fluc or pSG90 IVT RNA—individually or in combination—we introduced Mpa Ago along with a guide set designed to produce three distinct 72 nt fragments from Fluc and a single 100 nt fragment from pSG90 (Fig. S9A and S9B). Following cleavage, the reactions were analyzed by LC-MS/MS. In reactions containing only one RNA species, the corresponding target fragments from either Fluc (Fig. S9C) or pSG90 (Fig. S9D) were successfully enriched and identified over unassigned Ago-independent background fragments. Notably, in the reaction containing both RNA species, specific fragments derived from both Fluc and pSG90 were identified (Fig. S9E), indicating that Mpa Ago in combination with multiplexed guide sets enables precise RNA fingerprinting and thus identification of specific species in mixed RNA populations.

### Generation of custom RNA ladders using pAgos

The ability of pAgos to cleave a long RNA substrate into fragments of predetermined lengths defined by sequence-specific guides opens up the possibility of generating custom RNA ladders containing fragments of desired lengths. To this end, we synthesized a 1576 nt IVT RNA featuring six unique sequences of lengths ranging from 56 to 514 nt that were separated by a specific 20 nt sequence (5’-…CCUACCGUUAAUGACAGGUC… -3’, Fig. S10A). A single guide targeting this 20 nt region enabled simultaneous MpaAgo-mediated cleavage at sites between the adjacent sequences, releasing the six sequences from the parental RNA, thus generating a set of fragments with pre-defined lengths. Like in the reactions with multiplexed guides, the addition of the enhancer mix increased the cleavage efficiency and resulted in near complete cleavage of the substrate into a clean and precise RNA ladder, as shown on a denaturing 6% TBE-urea gel (Fig. S10B).

### RNA 5’ cap analysis using MpaAgo

The *N*7-methyl guanosine cap structure at the 5’ end of eukaryotic mRNAs is important for nuclear export, stability, and translation of mRNA *in situ*. Determining the efficiency of mRNA 5’ capping is an essential quality control step in modern mRNA vaccine/therapeutics manufacturing^43–45^. We sought to investigate whether pAgos could be favorably deployed for mRNA 5’ cap analysis. Capillary electrophoresis (CE) was used to assess cleavage efficiency and determine if pAgos could be programmed to cleave near the 5’ end of an RNA. A 5’-FAM labeled synthetic 33 nt oligoribonucleotide with a sequence representing the 5’ UTR sequence of Fluc mRNA was incubated with MpaAgo, and with one of four guides designed to induce cleavage 20 nt downstream of the 5’ end with either a complementary (g1A) or a non-complementary (g1C, g1G, g1T) 5’-most nucleotide (Fig. S11A). We found that MpaAgo efficiently and precisely cleaved at the intended site, regardless of the identity of the 5’ nucleotide, resulting in the expected 20 nt cleavage fragment (Fig. S11B).

Next, we extended this approach to full-length mRNAs. To facilitate capping efficiency testing in a straightforward and cost-effective gel-based assay, we designed guides that specifically targeted a site either 29 nt (Fluc mRNA) or 30 nt (pSG90 mRNA) downstream of their 5’ end (Fig. 3A). In both cases, the capped and un-capped cleavage products showed sufficient difference in mobility on a 15% denaturing polyacrylamide gel to observe successful capping (Figs. 3B and 3C).

**Figure 3.**
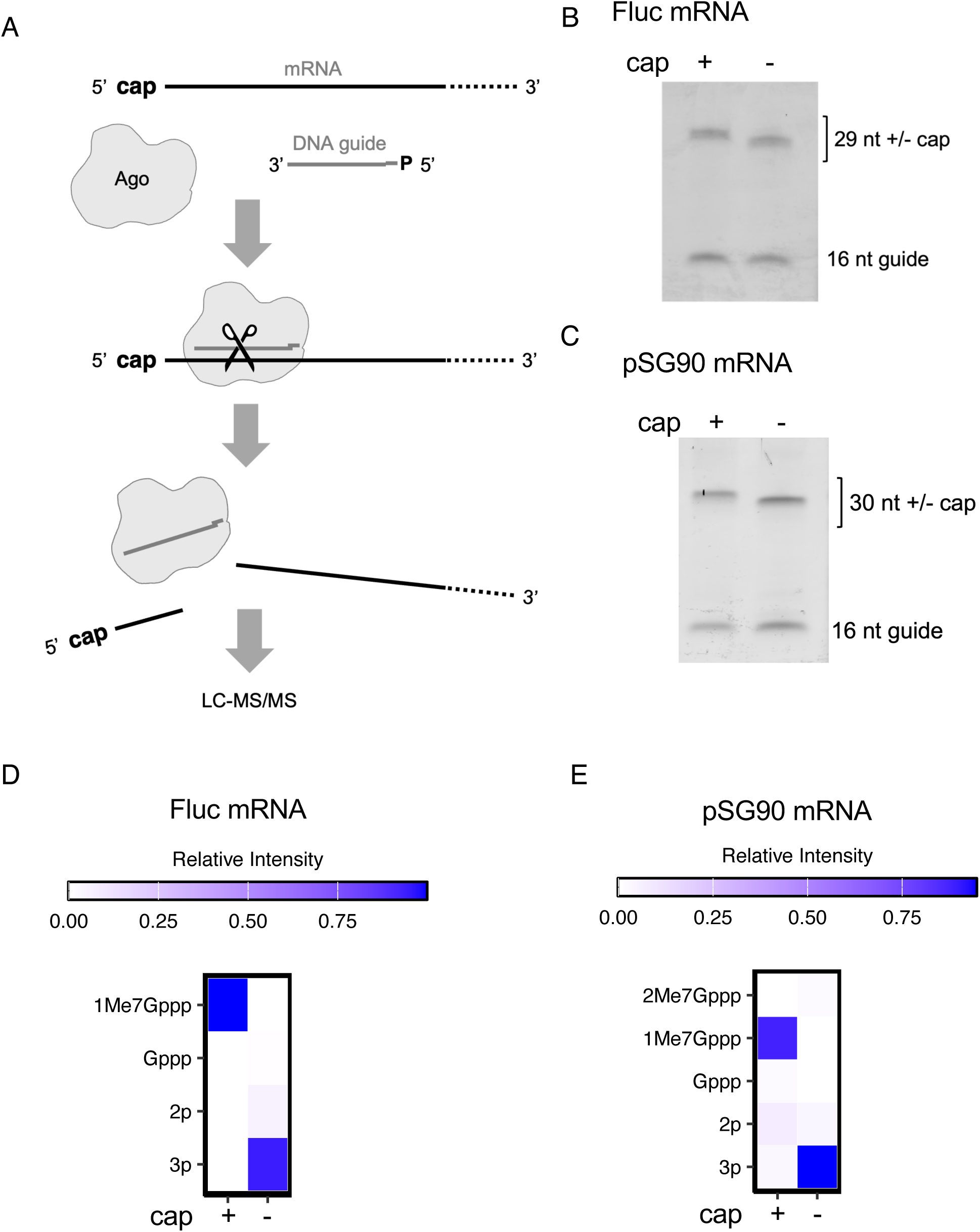
5’ cap analysis with MpaAgo. **(A)** Schematic of MpaAgo facilitated 5’ cap analysis, in which a 5’ phosphorylated guide targets MpaAgo cleavage to a site close to the 5’ end of a mRNA. Upon substrate release, the resulting fragments are analyzed via LC-MS/MS. **(B)** 5’ cleavage fragments (29 nt) of reactions containing Fluc mRNA that was either capped in vitro (+) or uncapped (-) were resolved on a 15% TBE-urea gel. Capped fragments migrate slower in the gel compared to uncapped fragments. The 16 nt guide DNA is indicated. **(C)** 5’ cleavage fragments (30 nt) of reactions containing pSG90 mRNA that was either capped in vitro (+) or uncapped (-) were resolved on a 15% TBE-urea gel. **(D)** and **(E)** Heatmap showing the relative intensity of detected 5’ fragments starting with a triphosphate (3p), diphosphate (2p), guanosine triphosphate (Gppp), cap 0 (1Me7Gppp), N⁷,2’-O-dimethylguanosine triphosphate cap (2Me7Gppp) for Fluc mRNA (D), and pSG90 mRNA (E).

To gain more detailed insight into the nature of the cap structure and estimate the capping efficiency, we turned to LC-MS/MS for the analysis of MpaAgo-mediated cleavage products. We targeted the 5’ Fluc-UTR using three distinct guides designed to generate 29 nt (guide Fluc_A1), 26 nt (guide Fluc_A1.2), and 32 nt (guide D891) 5’ fragments, respectively. We observed precise Fluc 5’ cleavage, producing fragments of the expected lengths 29-mer (90%), 26-mer (87%) and 32-mer (73%) (Fig. S12A). Some low levels of MpaAgo- and guide-independent 5 and 12 nt fragments were also detected (6-20%), which likely represent abortive transcripts (Fig. S12B). The cleavage of the pSG90 5’ UTR, too, was highly precise, predominantly yielding the intended 30 nt (D831 guide, >92%) or 21 nt (D892 guide, >94%) 5’ fragments (Fig. S12C). Similarly, MpaAgo- and guide-independent 5 and 12 nt fragments were the only other notable fragments detected. As expected, uncapped Fluc mRNA predominantly exhibited 5’ triphosphate termini (94%) and fewer diphosphate (5%) ends. In contrast, capped Fluc mRNA was composed exclusively of the Cap-0 (Me^7^Gppp) structure (Fig. 3D). Similarly, uncapped pSG90 mRNA consisted of 94% triphosphate and 4% diphosphate 5’ ends, whereas capped pSG90 mRNA comprised a mixture of 85% Cap-0, 7% diphosphate, and 3% triphosphate species (Fig. 3E).

These data show that MpaAgo can be used to successfully and quickly access the capping status and efficiency – either in a simple, cost-effective gel-based assay or by precise LC-MS/MS analysis.

### Homogenization of RNA 3’ ends and poly(A) tail analysis

Undesired 3’ extensions by RNA polymerases are a common byproduct of *in vitro* mRNA synthesis using T7 and other RNA polymerases and were first reported in the 1980s^46, 47^. Importantly, these 3’ extensions have been reported to negatively impact the efficacy of therapeutic mRNAs^48–50^.

We envisioned that MpaAgo could be employed to selectively target and cut a specific sequence incorporated into the poly(A) tail of an mRNA to remove unwanted extensions. Such 3’ end trimming would generate a mRNA population with homogenous 3’ ends (Fig. 4A). We utilized a slightly modified version of the poly(A) tail sequence utilized in the BNT162b2 vaccine^51^, consisting of a stretch of 30 adenosines, followed by a non-poly(A) sequence (GCATATGACTAAAAAACAT), and 60 terminal adenosines to a 610 nt IVT RNA sequence (pSG95). Guides D758-D761 were designed to induce MpaAgo cleavage in the non-poly(A) sequence (Fig. S13). After cleavage, the reaction products were resolved on a denaturing polyacrylamide gel. We found that guides D758, D760, and D761 enabled the complete cleavage of the full-length substrate at the respective target site, while a small fraction of the substrate remained uncleaved in the reaction with guide D759 (Fig. 4B). These data show that MpaAgo successfully homogenized the 3’ ends of IVT products, efficiently cleaving a target embedded in the poly(A) tail.

**Figure 4.**
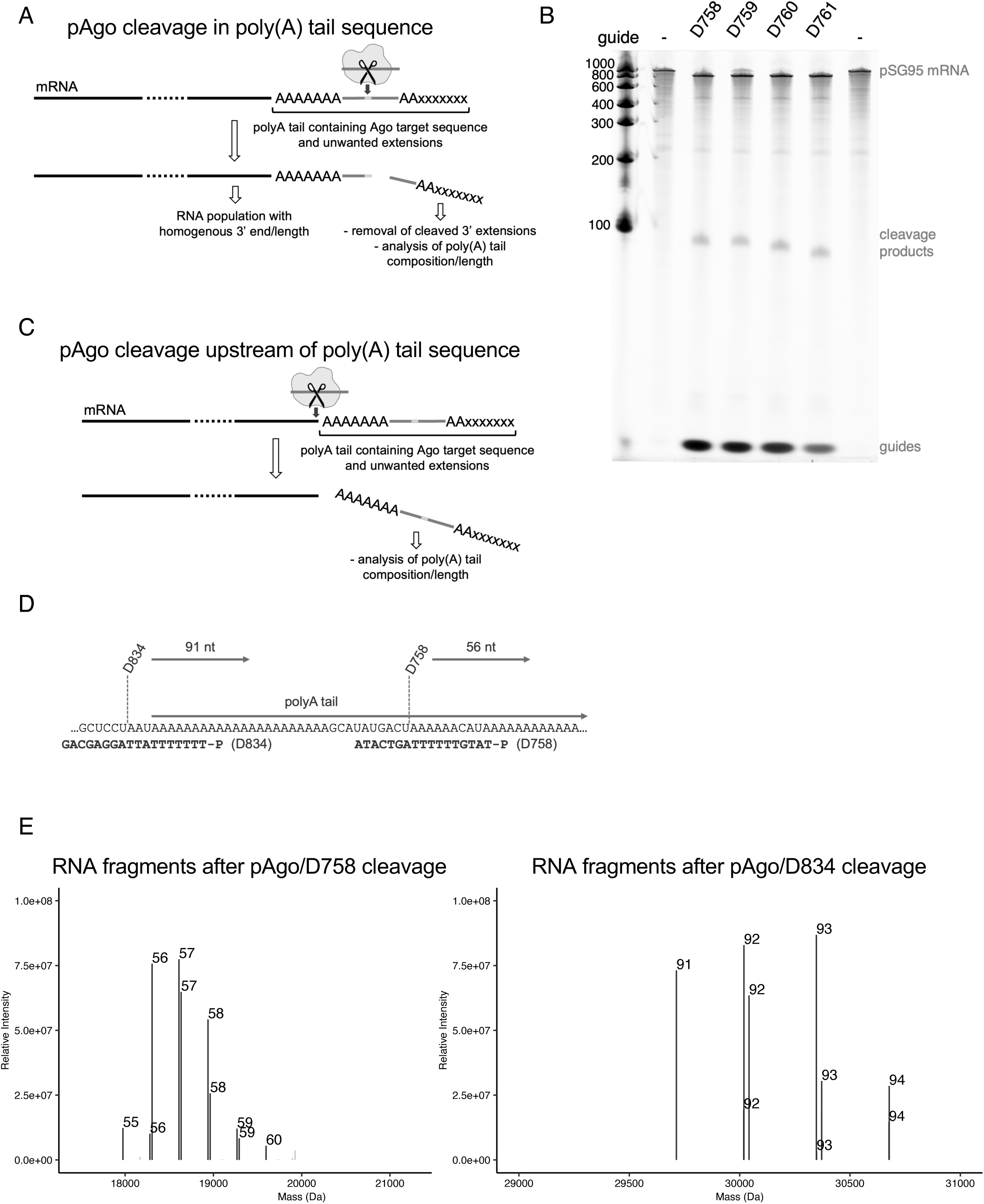
mRNA 3’ end processing by MpaAgo. **(A)** Schematic representation of mRNA 3’ end trimming. A non-poly(A) sequence island in the poly(A) tail of a mRNA is targeted and cleaved by MpaAgo, resulting in a mRNA population with homogenized 3’ ends, while the 3’ fragments can be further analyzed for their composition. **(B)** A 723 nt IVT RNA (pSG95) containing a non-poly(A) island was cleaved at four adjacent sites, using four guides (D758-D761 as indicated above the gel, Fig. S13), and the cleavage products were resolved on a denaturing 6% TBE-urea gel. **(C)** Schema of a reaction where MpaAgo is used to remove the complete poly(A) tail for analysis of the full poly(A) tail by LC-MS/MS. **(D)** Sequences of the 3’ region of the pSG120 IVT RNA and two guides (D758, D834). The respective cleavage sites and the lengths of the resulting 3’ fragments are indicated. **(E)** LC-MS/MS analysis of the 3’ fragments post cleavage using the D758 (left panel) and D834 (right panel) guides. The mass in kDa of the fragments is depicted on the x axis versus the relative intensity on the y axis. The lengths in nt are indicated above each bar.

Most current high-resolution methods to access poly(A) tail length and composition include the release of the poly(A) tail by RNase T1, which cleaves 3’ of guanosines^52^. However, when we attempted to use RNase T1 to measure poly(A) tail length of *in vitro* transcribed pSG120 mRNA, we found that RNase T1 also cleaved adenosines in a concentration-dependent matter (Fig. S14), necessitating optimization of RNase T1 concentration and digestion conditions for reliable poly(A) tail measurements. Here we show that for a more detailed analysis of poly(A) extensions and/or measurement of the poly(A) tail length using LC-MS/MS, MpaAgo can either be programmed to cleave directly upstream of the poly(A) tail (Fig. 4C) or a target sequence embedded in the tail as described above. The poly(A) tail in these experiments was slightly shortened to accommodate for the resolution of the LC-MS/MS, and to assure full capture of non-templated additions by T7 RNA polymerase without exceeding the range of the LC-MS/MS. MpaAgo was complemented with guides targeting a sequence upstream of the poly(A) tail (D834) or a sequence in the poly(A) tail (D758), resulting in 91 or 56 nt 3’ cleavage fragments, respectively (Fig. 4D). Cleavage reactions were performed as described above and the cleavage products analyzed by intact LC-MS/MS. Reactions with either guide allowed for the sequence determination of the 3’ cleavage fragments. As expected, in addition to the encoded poly(A) sequence, we observed multiple non-templated addition events by T7 RNA polymerase. In fact, we found that the majority of fragments had one or more additions to the encoded sequence (Fig. 4E). A significant fraction of fragments from both cleavage reactions were found to contain a single additional cytosine, an additional cytosine with one to three adenines, or one to three adenines without cytosine (Tables S1 and S2). These findings agree with what has been reported for non-templated T7 RNA polymerase additions^49^.

These data show that MpaAgo in combination with a guide targeting the poly(A) tail can be used to precisely cleave the 3’ end of mRNAs, thus creating mRNA molecules with homogenous ends. The 3’ cleavage fragments can be further analyzed by LC-MS/MS for poly(A) tail length measurement and assessment of non-templated additions by RNA polymerases.

## Discussion

Although the properties of DNA-targeting pAgos have long been acknowledged and utilized in biotechnological applications, RNA-targeting pAgos have largely been overlooked, likely due to their relative scarcity and limited characterization. In this study, we demonstrate that MpaAgo can serve as a powerful molecular tool, significantly enhancing or simplifying existing analytical workflows, such as RNA sequencing, fingerprinting, 5’ cap, and poly(A) tail analysis. Mpa Ago also enables the generation of mRNA populations with homogeneous mRNA 3’ ends.

We show that pAgos can be programmed with multiplexed guides in a single reaction to specifically and efficiently cleave at multiple sequences in a target mRNA. This suggests that pAgos may be used to address currently unresolved challenges in the analysis, quality control, and sequencing of long RNA molecules and therapeutics, such as self-amplifying RNAs (saRNAs), which exceed 10 kilobases in length. Current methods for mRNA analysis, such as digestion with sequence-specific endoribonucleases like RNase T1 or RNase 4, produce short RNA fragments (typically 10-20 nucleotides in length) that are subsequently analyzed by LC-MS/MS^53–56^. However, fragmenting mRNAs longer than 5 kilobases into such short fragments results in a complex mixture that is often challenging to deconvolute for full sequence reconstruction and/or modification detection. A possible solution to this problem is to increase the cleavage fragment size to a length that is short enough for mass spectrometry detection analysis, but long enough to reduce the complexity of the oligoribonucleotide pool for RNA sequencing by LC-MS/MS.

In our initial experiments aimed at targeting unique sites in Fluc IVT RNA using MpaAgo and multiplexed guides, the guides were designed to produce uniform cleavage fragments with a length of 72 nt. We found that addition of 0.5% BSA slightly increased the sequence coverage of the Fluc mRNA sequence, possibly by reducing non-specific binding of the loaded pAgo complexes to the RNA and other surfaces. Addition of an enhancer mix significantly increased the cleavage efficiency, and thus the controlled fragmentation of the RNA target, increasing the overall mapping coverage. We believe that the crowding agents in the enhancer mix facilitate the targeting of loaded pAgos to their RNA targets by increasing their proximity and thus their ability to interact in solution. Interestingly, adding the enhancer mix in combination with 0.5% BSA led to a reduction of coverage, albeit the gel analysis suggested more efficient cleavage. We hypothesize that the crowding agents increased non-specific binding of BSA to the RNA, potentially blocking access for MpaAgo to its target sites. Thus, BSA may be omitted for MpaAgo cleavage experiments that contain crowding agents (e.g., as part of the enhancer mix). Another factor potentially responsible for incomplete sequence coverage is the uniform length of the cleavage fragments, as fragments with the same length may not be identifiable/distinguishable due to identical or very similar masses.

Off-target cleavage of MpaAgo due to mismatched guide binding or slippage may also negatively affect fragment analysis. We thus set to design guides that a) were comprised of 18 nt instead of 16 nt to increase specificity, and b) induced cleavage into fragments of various lengths, sorted in size-bins ranging from 69 to 81 nt in length, to facilitate data deconvolution. While the band pattern on the gel suggested less efficient cleavage of the Fluc mRNA with the 18 nt guides (compare Fig. 2C, experiment 1 and Fig. S7A, experiment 2), the sequence coverage in reactions without the enhancer mix was comparable (∼ 79%). Reactions containing the enhancer mix reproducibly achieved slightly higher coverage (∼95%) using 18 nt guides than that of 16 nt guides (∼ 88%), suggesting that increasing guide specificity and cleavage of the substrate RNA into binned lengths may be beneficial for RNA sequence analysis. Upon closer inspection of the recovered fragments, we noticed that we were unable to detect fragments spanning Fluc mRNA positions 1056 to 1128 (16 nt guides) and 1064 to 1145 (18 nt guides) in any of our experiments. RNA structure modelling using MXfold2^57^ and RNAfold^58^ of the respective sequence and flanking regions did not indicate any obvious structural components that may inhibit pAgo binding (data not shown). The presence of the flanking fragments indicate that successful cleavage must have occurred. Sequencing of the IVT template ruled out the possibility of base mutations that could alter the mass of the expected fragments, making them unidentifiable by LC-MS/MS. We hypothesize that the missing fragments may be too similar in mass to other cleavage products that cannot be isolated in the data analysis. We are aware that the RNA sequences tested in this study are shorter than those of saRNAs, and that the maximal number of combined guides tested was 24. However, our results suggest that this application could successfully be used with clinically relevant RNA substrates, especially since full replacement of uridine with m^1^Ψ, a replacement often found in therapeutic mRNAs to minimize the host immune response, only had a moderate effect on the sequence coverage.

pAgos in combination with a smaller pool of multiplexed guides that cleave RNA substrates to generate a unique, characteristic, and identifiable fragment pattern can be used to fingerprint RNAs. This may be useful to confirm the presence of certain RNAs in an RNA mixture (e.g., to analyze the RNA molecule distribution in RNA pools packaged in liquid nanoparticles) or quick detection of a specific target RNA in a biological sample for diagnostic workflows. A similar approach was recently developed using the HD3 deoxyribozyme to fragment RNA^59^. Due to its single turnover character, this method requires multiple rounds of cleavage and re-annealing of the HD3 deoxyribozyme pool, which requires up to 24 h total digestion time for complex substrates. This is much slower than the 15 min of cleavage time in the MpaAgo reaction. In addition, the target site choice is limited to a particular set of trinucleotide motifs at which HD3 cleaves, a limitation that does not apply to MpaAgo cleavage.

In eukaryotic mRNA, the *N*7-methyl guanosine cap structure is a 5’ end modification that is important for nuclear export, RNA stability, and translation. Thus, effective capping of mRNA is an essential part of mRNA-based vaccine and therapeutic synthesis. Current methods of cap analysis combine LC-MS/MS analysis of a defined 5’ end fragment of a mRNA that was generated by selective cleavage using either *trans* acting deoxyribozymes^60–62^, and ribozymes^63^, or a DNA probe-directed approach using either the sequence-nonspecific Ribonuclease H (RNase H)^64^, or sequence specific RNases (e.g., RNase 4, MC1, and RNase T1^65, 66^) to cleave the 5’ UTR at a specific position and release the cap-containing 5’ fragment.

Our data show that guided MpaAgo can efficiently release the cap-containing 5’ termini of mRNAs, enabling the analysis of mRNA capping efficiency and permitting the determination of the cap structure comparably to other approaches, such as RNase H or sequence-specific RNases. Contrary to non-guided endoribonucleases, pAgos exhibit lower turnover^23, 30^, thereby requiring comparatively high enzyme concentrations for effective activity. However, there are significant advantages of using MpaAgo for cap analysis. First, it eliminates the need for a blocking antisense oligo or the presence/incorporation of an endoribonuclease recognition sequence for precise cleavage. Successful 5’ fragment generation by sequence-specific endoribonucleases (e.g., RNase T1, RNase 4) requires blocking of all the respective target sites directly downstream of the 5’ end by hybridization of a DNA oligo. In addition, the release of the protected fragment is dependent on the presence of a recognition site directly downstream of the duplex. Second, sequence-specific RNases will fragment the entire mRNA downstream of the hybridized probe, in some workflows necessitating the enrichment of the 5’ cleavage fragments. Moreover, the additional step of hybridizing of a blocking oligo is not needed with MpaAgo. The use of MpaAgo for 5’ end analysis may also allow for the detection of endogenous 5’ ends with poorly defined transcription start sites or post-transcriptional modifications^67^, in which case the use of a blocking oligo may not be sufficient to protect the heterogeneous 5’ ends. Additionally, the transient alignment of the short DNA guide with the RNA target at 50°C may mitigate the influence of RNA structure in the 5’ UTR sequence, reducing bias and increasing binding/cleavage efficiency in MpaAgo workflows.

A potential caveat of cap analysis with sequence-specific RNases is that it generates RNA/DNA duplexes with a 3’-phosphate and/or 2’,3’-cyclic-phosphate overhang. In contrast, the release of single-stranded MpaAgo cleavage products facilitates subsequent 3’ end modifications of the fragments, such as fluorescent labeling through fill-in extension using DNA polymerases as in RNase H-based cap analysis methods^64^, or enzymatic ligation of the fragments to other nucleic acid sequences.

In addition to requiring a significantly shorter reaction time compared to other workflows (15 min vs. 60 min), MpaAgo also facilitates 5’ cap analysis by producing cleavage products that do not need to be enriched for LC-MS/MS analysis. In fact, the generation of only two cleavage products (a short 5’ fragment and a long 3’ fragment) allows for an easy and cheap capping efficiency assessment using PAGE, making this analysis accessible to laboratories without LC-MS/MS capabilities. Recently, a group reported the use of a nucleolytic toxin from a type III toxin/antitoxin system that recognizes a specific 5 nt target sequence to cleave and subsequently analyze 5’ mRNA ends for cap detection^68^. Theoretically, mRNA cleavage with Toxin III endoribonucleases also generates only two fragments that enable gel-based capping assessment, and they do not require a DNA guide or oligo for sequence specific cleavage. However, this approach necessitates sequence optimization of the mRNA to remove other potential Toxin III cleavage sites, and – more importantly - a change of the 5’ UTR sequence to incorporate the target sequence, which may be limiting for certain mRNA constructs. In addition, type III toxins possess a relative specificity rather than an absolute specificity, meaning that they are prone to off-target cleavage at similar sequence motifs^69^. This can result in the generation of additional fragments, which may interfere with cap analysis.

One of the shortcomings of using RNase H for cap analysis is its heterogeneous cleavage that results in additional fragments at positions directly upstream or downstream of the desired cleavage site. Optimization of the DNA oligo sequence is often required to reduce off-target cleavage^64^. We show that there is only negligible off-target cleavage by MpaAgo as tested with four different guides on two different mRNAs by CE and LC-MS/MS.

We also show that site-directed cleavage by MpaAgo is a superior tool when analyzing RNA 3’ ends. The 3’ poly(A) tail directly affects mRNA stability and translation efficiency, making it an essential component of native and therapeutic mRNAs. While several low-resolution methods exist that can identify the presence of a poly(A) tail, there are only limited options for detailed analysis of the poly(A) tail length and composition. In addition to next generation sequencing, other high-resolution methods include digestion of the RNA substrate with RNase T1, which cleaves after every guanine, followed by capillary gel electrophoresis or LC-MS/MS^70, 71^. Site-specific removal of the poly(A) tail by MpaAgo for subsequent analysis addresses several shortcomings of digestion with RNase T1. Modern mRNA therapeutics and vaccines incorporate G-containing, non-A islands into the poly(A) tails to increase RNA polymerase accuracy (e.g., Pfizer BNT162b2). Guanines in these sequences will be targeted by RNase T1 and generate a diverse pool of poly(A) fragments which complicates analysis. In addition, we show here that RNase T1 can non-specifically degrade poly(A) regions in a concentration dependent manner. MpaAgo on the other hand can be directed to precisely cleave either the complete poly(A) tail or fragments thereof when targeting the non-A islands. This generates defined cleavage products, which abolishes the need for subsequent purification or enrichment. Furthermore, it simplifies the subsequent analysis by enabling noise-free, single-nucleotide resolution analysis of the poly(A) tail via LC-MS/MS.

In addition, MpaAgo uniquely enables 3’ end polishing of IVT RNA products to generate RNA populations with homogenous 3’ ends. Non-templated addition of nucleotides is a known feature of RNA polymerases *in vitro* that can lead to unwanted consequences (e.g., providing a source for double-stranded RNA (dsRNA) formation). Targeted trimming and thus homogenizing of 3’ RNA ends with MpaAgo may prevent and/or significantly reduce the formation of immunogenic dsRNA in RNA manufacturing, addressing a main concern in RNA therapeutics development. Since the MpaAgo enzyme, the short DNA guide, and the cleaved RNA extension can easily be removed from the homogenous RNA pool by proteinase K/DNase I treatment, and/or a sizing column, this workflow could be incorporated into large scale RNA synthesis workflows.

Taken together, we show that programmed cleavage of mRNA by MpaAgo enables faster, more precise, and straightforward workflows for multiple important RNA analysis workflows, like 5’ capping analysis, RNA sequence/modification mapping, and poly(A) tail analysis – all essential for therapeutic RNA quality assessment. In addition, we show that MpaAgo can be used to generate RNA populations with homogeneous 3’ ends, which may help to minimize the presence of immunogenic dsRNA and thus address one of the most critical hurdles in therapeutic RNA production.

## Methods

### *In vitro* RNA cleavage assay

200 ng of Fluc RNA (∼1.8 kb, ∼0.37 pmol), a shortened version of the pTsin saRNA (pSG90; ∼1.7 kb, ∼0.37 pmol), or Epo RNA with a modified sequence (pSG95; ∼0.7 kb, ∼0.86 pmol), all synthesized using *in vitro* transcription (IVT) (New England Biolabs, #E2040) according to the manufacturer’s instructions, were cleaved with MpaAgo in a 10 µL reaction at 50°C for either 15 minutes (single and double cleavages, 1:5 substrate: Ago molar ratio) or 30 minutes (multiplexing, 1:10 substrate: Ago ratio) in 1x ThermoPol® reaction buffer (New England Biolabs #B9004S). The Ago:guide molar ratios were 1:5 for single and double cleavages, 1:1 for multiplexing experiments, and 1:10 for cleaving polycistronic RNA. DNA guides were pooled using the Echo acoustic liquid handler (Beckman Coulter) and phosphorylated using T4 polynucleotide kinase (New England Biolabs #M0201S) according to the manufacturer’s protocols. For multiplexing experiments and when indicated, a 1x enhancer mix containing 0.5 M betaine, 20 mM potassium glutamate, 20% glycerol, and 5% PEG3350 was added to the reactions. BSA was added to 0.5% (v/v) when indicated. Removal of DNA guides was performed by adding 20 U of exonuclease I (New England Biolabs, #M0293), followed by incubation at 37°C for 15 minutes. Cleavage experiments were stopped by the addition of 0.08 U of proteinase K (New England Biolabs #P8107S) at room temperature for 5-10 minutes. After adding RNA loading dye (New England Biolabs, #B0363), cleaved RNAs were resolved by gel electrophoresis on 6% TBE-urea gels (Thermo Fisher Scientific #EC6265), stained by 0.5x SYBR Gold (Thermo Fisher Scientific #S11494), and analyzed using the Typhoon biomolecule imager (Cytiva) or Odyssey M imager (Licor Bio). For mass spectrometry analysis, reactions were scaled up 10-fold in ThermoPol® reaction buffer (i.e., 2 µg of RNA) at 50°C for 30 minutes. Cleavage products were then column-purified (New England Biolabs #T2030S) and eluted in 30 µL of nuclease-free water. Purified RNA was subsequently filtered with Ultrafree-MC Centrifugal Filter (Millipore Sigma # UFC30GV0S) for 5 min at 10k x g prior to LC-MS/MS analysis. For RNA fingerprinting, cleavage assays were performed as described above, but with a limited set of guides targeting Fluc IVT RNA and pSG90 IVT RNA, generating three 72 nt or one 100 nt RNA species-specific cleavage fragments, respectively. 3.5 pmol of Fluc IVT RNA, 3.5 pmol of pSG90 IVT RNA, or a mix of 1.75 pmol Fluc IVT RNA and 1.75 pmol pSG90 IVT RNA were incubated with 17.5 pmol of the guide pool in the presence or absence of 17.5 pmol MpaAgo in 1x ThermoPol® reaction buffer and the enhancer mix for 30 minutes at 50°C. Reactions were stopped by the addition of 0.8 units proteinase K and incubation at 24°C for 10 minutes. Following a column cleanup, samples were either ran on a denaturing 6% TBE-urea gel or analyzed by LC-MS/MS.

### Capping analysis

For 5’ mRNA cap analysis experiments, 50 µg (∼ 83 pmol) or 25 µg (∼ 44 pmol) of Fluc or pSG90 *in vitro* transcribed mRNA, respectively, were initially denatured for 5 min at 65°C and immediately put on ice. Both mRNAs were capped using FCE (New England Biolabs #M2081S) according to the manufacturer’s manual. The capping reactions were subsequently column purified (New England Biolabs #T2040S). 200 µl reactions containing 3.5 pmol of capped and uncapped Fluc or pSG90 mRNA were then incubated for 15 min at 50°C in 1x ThermoPol® reaction buffer, 30% glycerol, 160 units of Murine RNase Inhibitor (New England Biolabs #M0314S), 67.4 pmol MpaAgo, and 66.4 pmol guide (A1: /5Phos/ATCTCCTTCTTAAAGT, A1.2: /5Phos/ TCCTTCTTAAAGTTAAAC, D891: /5Phos/ TATATCTCCTTCTTAA targeting Fluc mRNA, or D831: /5Phos/GTCGGCTGTTTGATTC, D892: /5Phos/ TTGATTCAATAGTGTG targeting pSG90 mRNA). Reactions were incubated at 50°C. After 15 min, the reactions were put on ice, 120 units of Exonuclease I (New England Biolabs #M0293S) was added, and the reactions incubated at 37°C for 15 min to remove unbound DNA guides. The reactions were stopped by the addition of EDTA to 20 mM and incubation at room temperature for 5 min. Finally, reactions were column purified and submitted for LC-MS/MS analysis.

### 3’ end homogenization and poly(A) tail analysis

For the analysis of the poly(A) tail composition and length and to homogenize the 3’ ends of a mRNA population, MpaAgo reactions were scaled up to a 400 µl reaction volume. 67 pmol of *in vitro* transcribed pSG95 (723 nt) or pSG120 (698 nt) mRNAs were incubated with 162 pmol MpaAgo, 800 pmol D758 (/5Phos/TATGTTTTTTAGTCATA) or D834 (/5Phos/TTTTTTTATTAGGAGCAG) guides in 1x ThermoPol® reaction buffer with 640 units of Murine RNase Inhibitor for 30 min at 50°C. Reactions were stopped by the addition of 1.6 units proteinase K and 10 min incubation at room temperature. Reactions were column purified, followed by enrichment of the poly(A) containing cleavage fragments. The volume of the purified cleavage fragments was adjusted to 50 µl, then 1 volume (50 µl) of high salt buffer (20 mM Tris/Cl pH 7.5, 0.5 M LiCl_2_, 10 mM EDTA) was added. 250 µl of oligo d(T)_25_ magnetic beads (New England Biolabs #S1419S) were washed once with 100 µl low salt buffer (20 mM Tris/Cl pH 7.5, 50 mM LiCl_2_, 10 mM EDTA), and twice with high salt buffer. The RNA was added to the beads and incubated at room temperature on a rotator for 20 min. Bead-bound RNA was washed once with low salt buffer, twice with high salt buffer, and one final time with low salt buffer. To elute the enriched RNA, the beads were incubated with 30 µl of H_2_O for 2 min at 50°C in a heat shaker (400 rpm). The eluted RNA was recovered, incubated for 5 min at 65°C, and spun through Ultrafree-MC Centrifugal Filter (Millipore Sigma # UFC30GV0S) for 5 min at 10k x g. The enriched and filtered samples were then subjected to LC-MS/MS analysis.

## Acknowledgements

The authors want to thank Katherine Marks for purifying the pAgo variants tested in this study. We also want to thank Thomas C. Evans Jr. and Eric Cantor for their advice and strong support of this research.

## Ethics declarations

At the time of this work, the authors were employed by New England Biolabs, Inc., which funded this work and is the manufacturer of reagents, some of which were used in this manuscript. Even though the authors were employed and funded by New England Biolabs, Inc., this did not detract from the objectivity of data generation or its interpretation.

