## Supplemental Material for "Multifaceted mRNA analysis using programmed RNA cleavage by *Mucilaginibacter paludis* Argonaute"

#### Supplemental Methods

##### Cleavage of polycistronic RNA

Targeting multiple sites with a singular guide can be used to generate custom RNA ladders. We inserted five D770 guide target sequences (/5Phos/ NCTACCGTTAATGACAGGTC) into a 1576 nt *in vitro* transcribed RNA (pSG106 RNA) so that successful cleavage by MpaAgo loaded with the D770 guide would result in the generation of six RNA fragments with various lengths from 56 to 514 nt. 0.37 pmol of the pSG106 RNA were incubated with 3.7 pmol of MpaAgo and 18.7 pmol of guide 770 in either 1x ThermoPol®, 1x NEB r1.1, or T4 DNA ligase reaction buffer as indicated, Murine RNase inhibitor without or with crowding agent (30% glycerol) for 30 min at 50°C. The reactions were stopped by the addition of proteinase K and incubation at 37°C for 10 min. After the addition of RNA loading dye and heat denaturation (5 min at 70°C), the fragments were resolved by denaturing gel electrophoresis.

##### Poly(A) tail analysis using RNase T1

Poly(A) analysis with RNase T1 was performed for a pSG120 IVT mRNA substrate with an encoded poly(A) tail. Briefly, 10 $\mu$g of pSG120 mRNA was incubated with 500 – 4000 U of RNase T1 (Thermo Cat # ENO541) in 50 mM Tris HCl, pH 7.5; 2 mM EDTA to a final reaction volume of 30 $\mu$L. Each reaction was incubated at 37°C for 1 hour. Following incubation, each reaction was diluted to 50 $\mu$L in water and combined with 50 $\mu$L of high salt buffer (20 mM Tris-HCl pH 7.5, 500 mM LiCl2, 10 mM EDTA). Each reaction was combined with 100 $\mu$L of washed Oligo d(T)$_{25}$ Magnetic Beads (NEB Cat # S1419S) and incubated at room temperature for 20 minutes with occasional agitation to complex poly(A) species with the Oligo d(T)$_{25}$ beads. Next, complexed Oligo d(T)$_{25}$ beads were washed with once low salt buffer (20 mM Tris-HCl pH 7.5, 50 mM LiCl2, 10 mM EDTA), twice with high salt buffer, and once with low salt buffer. Complexed poly(A) species were eluted from the beads by heating to 65°C in water for 5 minutes. Elutions were spin-filtered with UltraFree-MC/GV centrifugal filters at 10K x g for 5 minutes and analyzed by UHPLC-MS/MS.

**Oligonucleotide UHPLC-MS/MS**

Oligonucleotide separation was performed on a Vanquish Horizon UHPLC equipped with a Waters Acquity Premier BEH C18 130A, 1.7 $\mu$m 2.1 x 100mm reverse phase column, heated to 70°C. A 20-minute gradient of solvent A (1% hexafluoroisopropanol (HFIP), 0.1% diisopropylethylamine (DIEA), 1 $\mu$M EDTA in water) and solvent B (90% Methanol, 0.075% HFIP, 0.0375% DIEA, 1 $\mu$M EDTA) was applied at a flow rate of 300 $\mu$L/min for oligonucleotide separation. High-resolution RNA oligonucleotide MS$^1$ data was acquired on an Orbitrap Eclipse mass spectrometer. MS$^1$ data was acquired at a resolution of 120,000 (for Poly(A) and cap analysis) and 240,000 (for fingerprinting analysis). Deconvoluted RNA oligonucleotide masses were calculated using Thermo BioPharmaFinder 5.1 software. Sequence annotation of observed masses was performed utilizing in-house scripts.

#### Supplemental Figure Legends

**Figure S1. Site-specific Mpa Ago cleavage of IVT RNA**

pSG95 IVT RNA (~0.7 kb), a U-depleted version of human Epo mRNA with modified sequences and a high GC-content (~59%), was subject to MpaAgo cleavage under the same conditions as shown in Fig. 1. 5' phosphorylated 16 nt guides were designed to target sites 120 nt apart. Cleavage products were analyzed on a denaturing 6% TBE-urea gel stained with SYBR Gold. Guide identities and cleavage efficiencies (%) were indicated above the corresponding lanes and the uncut full-length IVT RNA (pSG95 RNA), guides (16 nt), and other cleavage fragments are indicated on the side of the gel.

**Supplemental material**

**Figure S2. Mpa Ago-mediated RNA cleavage is guide-dependent.**
pSG90 IVT RNA (~1.7 kb) was subject to MpaAgo cleavage in ThermoPol® reaction buffer at 50˚C for 15 minutes with single 5' phosphorylated DNA guides that were not complementary to pSG90 RNA. The substrate:Ago molar ratio was 1:5, and the Ago:guide molar ratio was 1:5. Cleavage products were analyzed on a denaturing 6% TBE-urea gel stained with SYBR Gold and no guide-facilitated cleavage was detected.

**Figure S3. Cleavage of Fluc IVT RNA by MpaAgo with 18 nt guides.**
Fluc IVT RNA (~1.8 kb) was subjected to MpaAgo cleavage with single complementary 5' phosphorylated DNA guides in the ThermoPol® reaction buffer at 50˚C for 15 minutes. The substrate:Ago molar ratio was 1:5, and the Ago:guide molar ratio was 1:5. The 18 nt guides were designed generate cleavage fragments between 69 to 81 nt in length to aid with data analysis. Cleavage products were analyzed on a denaturing 6% TBE-urea gel stained with SYBR Gold. Guide identities and cleavage efficiencies (%) were indicated above the corresponding lanes.

**Figure S4. MpaAgo is active at various temperatures.**
**(A)** pSG90 IVT RNA (~1.7 kb) was subject to MpaAgo cleavage with a single complementary DNA guide in ThermoPol® reaction buffer at 4˚C, 24˚C, 37˚C, or 50˚C for 10, 15, 30, 60, or 240 minutes. The substrate:Ago molar ratio was 1:5, and the Ago:guide molar ratio was 1:5. Cleavage products were analyzed on a denaturing 6% TBE-urea gel stained with SYBR Gold. The full length, uncut pSG90 mRNA band is labeled with a grey asterisk, the cleavage products are indicated by a bracket. The guide band is labeled with two asterisks. **(B)** pSG90 IVT RNA (grey asterisk) was incubated with or without MpaAgo/guide at 4°C for 16 h, and at 24°C/37°C/50°C for 4 h as indicated above the lanes. The guide band is labeled with two asterisks.

**Figure S5. DNA guides can be efficiently removed from pAgo reactions.**

**Supplemental material**

pSG90 IVT RNA (~1.7 kb) was subject to MpaAgo cleavage with a complementary DNA guide in the ThermoPol® reaction buffer at 50˚C for 15 minutes. The substrate:Ago molar ratio was 1:5, and the Ago:guide molar ratio was 1:5. After MpaAgo cleavage, *E. coli* exonuclease I was added directly to the reactions to evaluate its ability to remove DNA guides.

**Figure S6. Fluc sequence coverage determined by LC-MS/MS.**
Graphical representations of Fluc sequence coverage data shown in Fig. 2D. Experimental conditions: 1. no guide control, 2. pool of 24 guides, no additives; 3. pool of 24 guides with 0.5% BSA + enhancer mix; 4. pool of 24 guides with 0.5% BSA; and 5. pool of 24 guides with the enhancer mix. The black bars represent the detected RNA fragments along the complete Fluc RNA, and the blue bars represent the abundance of the respective fragments. The percent complete sequence coverage for each experiment are indicated.

**Figure S7. Cleavage of RNAs into fragments of various sizes further improved sequence coverage determined by LC-MS/MS.**
**(A)** Fluc IVT RNA (~1.8 kb, experiments 1-3) or pSG90 IVT RNA (~1.7 kb, experiments 4-6) were subjected to MpaAgo cleavage with or without a pool of 22 or 21 5' phosphorylated guides, respectively, and with or without enhancer mix as indicated, at 50˚C for 30 minutes. The substrate:Ago molar ratio was 1:10, and the Ago:guide molar ratio was 1:1. The guides were designed to produce cleavage fragments of sizes between 69-81 nt when multiplexed as indicated by the bracket next to the gel. Cleavage products were analyzed on a denaturing 6% TBE-urea gel stained with SYBR Gold. **(B)** Cleavage products of two biological replicates of the experiment shown in (A) analyzed by LC-MS/MS to determine the sequence coverage for either Fluc (experiments 1-3) or pSG90 (experiments 4-6) RNA . Sequence coverages (%) are shown with standard error bars. **(C)** Graphical representations of Fluc sequence coverage data of one replica of the reactions shown in (B) (Top: no enhancer mix, experiment 2; bottom: with enhancer mix, experiment 3). The black bars represent the

detected RNA fragments along the complete Fluc RNA. The blue bars show the abundance of the respective fragments. **(D)** Same as (C), but for the reactions containing pSG90 RNA.

**Figure S8. MpaAgo cleaves fully m1Ψ-modified Fluc mRNA with reduced efficiency.**

**(A)** Fully m1Ψ-modified Fluc RNA was subject to MpaAgo cleavage with or without a pool of 22 5' phosphorylated 18 nt guides, and with or without enhancer mix as indicated above the lanes. The substrate:Ago molar ratio was 1:10, and the Ago:guide molar ratio was 1:1. Cleavage products were analyzed on a denaturing 6% TBE-urea gel stained with SYBR Gold. Experiments 4 and 5 are replicates using RNA substrates synthesized in independent reactions. **(B)** Sequence coverage in percent derived from the Fluc cleavage products from (A) as determined by LC-MS/MS. **(C)** Graphical representations of m1Ψ-modified Fluc sequence coverage data (Top: no enhancers; bottom: with enhancers) shown in (B). Detected RNA fragments are represented by black bars along the complete Fluc RNA. The blue bars show the abundance of the respective fragments.

**Figure S9. Mpa Ago enables RNA fingerprinting**

**(A)** Schematic representation of an RNA fingerprinting experiment, where an equimolar mix of Fluc and pSG90 IVT RNA substrates were incubated with Mpa Ago and guides that were designed to generate three specific 72 nt cleavage fragments of Fluc (fr. 1-3) and one 100 nt cleavage fragment of pSG90 (fr. 4) RNA. **(B)** Gel-fractionated products of experiments described in (A), in which either Fluc (F; lanes 1 and 3) or pSG90 (90; lanes 2 and 4) IVT RNA or a combination of both (lane 5) was incubated with a guide pool that in the presence of MpaAgo resulted in the formation of 72 nt fragments for Fluc and a 100 nt fragment for pSG90. **(C)** LC-MS/MS analysis of the specific Fluc or **(D)** pSG90 fragments post cleavage using the guide pool. A no-enzyme control is shown in the top panel. The mass in kDa of the fragments is depicted on the x axis versus the relative intensity on the y axis. The name of the identified fragment (salmon) is indicated above each bar. Unidentified background fragments are shown in cyan. **(E)** LC-MS/MS analysis of cleavage fragments from a reaction containing both, Fluc and pSG90 IVT RNA substrates.

**Figure S10. Generation of custom RNA ladders using Mpa Ago.**
**(A)** Schematic representation of the pSG106 RNA (1576 nt), which contains six RNA sequences with lengths from 56 to 514 nt that are separated by the recognition sequence for a 5' phosphorylated DNA guide. The location of the guide recognition sequence is indicated by asterisks. **(B)** pSG106 RNA was subjected to cleavage with MpaAgo and the 5' phosphorylated D770 guide for 30 min at 50°C in ThermoPol® reaction buffer at 1:10 substrate:MpaAgo and 1:5 MpaAgo:guide ratios with or without addition of the enhancer mix as indicated above the lanes. Bands showing successfully cleaved fragments 1-6 are labeled and the uncut, full length pSG106 RNA is marked with a black asterisk.

**Figure S11. Mpa Ago precisely cleaves the 5' end of RNA.**
**(A)** Schematic representation of a cleavage reaction, in which Mpa Ago is guided to cleave close to the 5' end of a 5'-FAM (green star) labeled RNA, generating a 20 nt 5' fragment and a 13 nt 3' fragment. **(B)** A 5' FAM labeled RNA substrate (SynFLuc8.1, 33 nt) was designed to mimic the native 5' UTR Fluc sequence. Complementary guides were designed with all possible 5' position 1 bases (g1x). Mpa Ago in combination with the guides were used to cleave the RNA and the cleavage products were analyzed via capillary electrophoresis. The fragment size was confirmed with a positive control substrate of the same size (not shown). The 5' cleavage product is marked with a dotted line and annotated with its size, while the uncleaved substrate is marked with an asterisk. All 5' cleavage products were the expected size, and the lack of peak shoulders indicate precise cleavage rather than slippage at the 10-11 cut site. RFU values vary with capillary injection conditions and are not indicative of reaction completeness.

**Supplemental material**

**Figure S12. LC-MS/MS detection of 5' cleavage products.**

**(A)** Heatmap showing the relative intensity and the lengths of detected 5' fragments after MpaAgo cleavage of capped Fluc mRNA using three different DNA guides (A1: /5Phos/ATCTCCTTCTTAAAGT (29 nt expected cleavage fragment length), A1.2: /5Phos/ TCCTTCTTAAAGTTAAAC (26 nt expected fragment length), D891: /5Phos/ TATATCTCCTTCTTAA (32 nt expected fragment length). **(B)** Control reactions in which capped Fluc mRNA was incubated with the D891 guide but without MpaAgo or with MpaAgo, but without any guide are shown in the right panel. **(C)** Length distribution of capped 5' pSG90 mRNA fragments after MpaAgo cleavage, showing two controls (1. guide, but no MpaAgo, 2. MpaAgo, no guide) and MpaAgo reactions with guides (D831: /5Phos/GTCGGCTGTTTGATTC (30 nt expected fragment length), D892: /5Phos/ TTGATTCAATAGTGTG (21 nt expected fragment length). The 5' UTR sequences of the respective mRNA substrates and the predicted 5' cleavage fragments are shown below the heatmaps.

**Figure S13. MpaAgo facilitates poly(A) tail analysis.**

The 3' terminal sequence of the pSG120 mRNA with parts of the poly(A) tail are shown. The target sequences of 5' phosphorylated DNA guides used in experiments shown in Fig. 4 and their respective cleavage sites are indicated.

**Figure S14. Poly(A) tail analysis with RNase T1**

**(A)** UV chromatogram of oligo-DT isolated species from pSG120 mRNA following digestion with varying concentrations of RNase T1. The chromatographic peaks of full-length RNase T1 cleavage products (5'-Poly(A) & 3'-Poly(A))and those possibly resulting from poly(A) cleavage (Cleaved Poly(A) Species) events are annotated. **(B)** A bar graph of the summed relative intensity of full-length RNase T1 digestion products (Full-Length), shortened poly(A) products (Cleaved) and unidentified masses from oligo-DT isolations of pSG120 mRNA following digestion with varying concentrations of RNase T1.

**Supplemental material**

**Table S1.** LC-MS/MS data used to generate Fig. 4E (D758 guide for cleavage in a non-poly(A) island in the poly(A) tail), indicating the experimental and theoretical masses, as well as the sequences of the detected RNA fragments.

**Table S2.** LC-MS/MS data used to generate Fig. 4E (D834 guide for removal of the complete poly(A) tail), indicating the experimental and theoretical masses, as well as the sequences of the detected RNA fragments.

**Table S3.** Sequences of mRNAs used in this study

Supplemental Figure 1

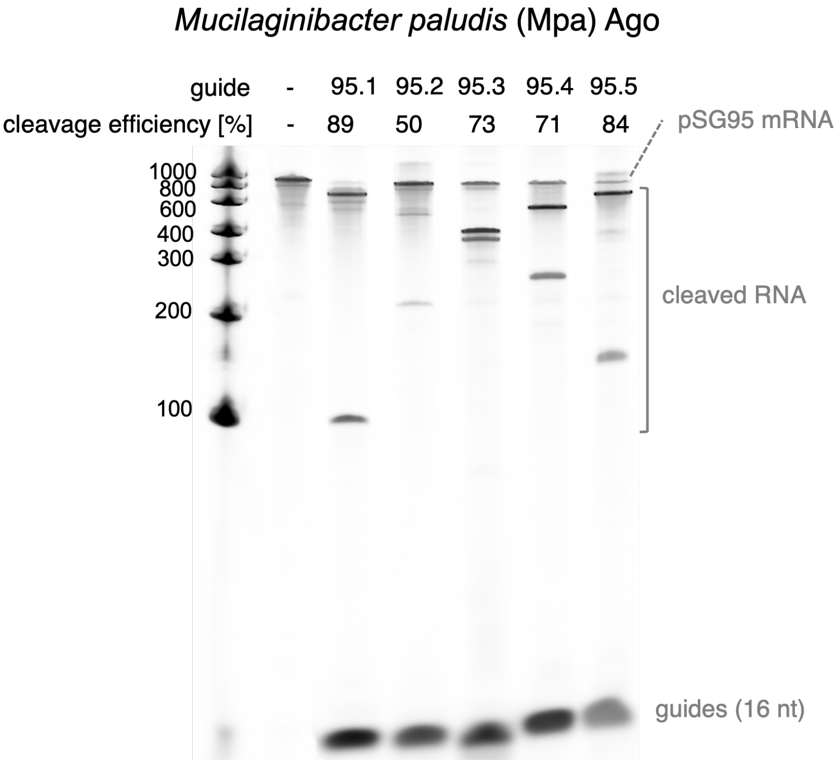

Supplemental Figure 2

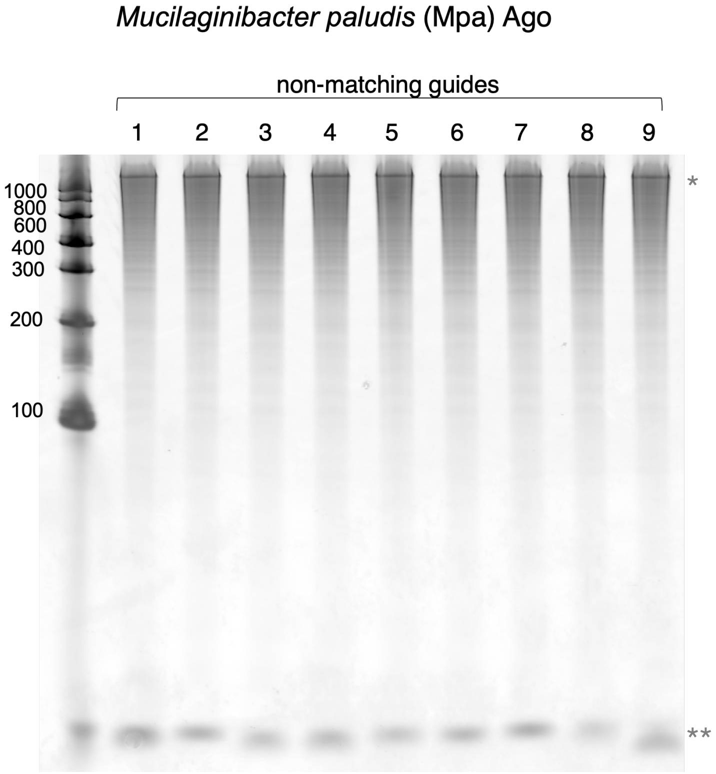


*pSG90 IVT RNA
**guides

Supplemental Figure 3

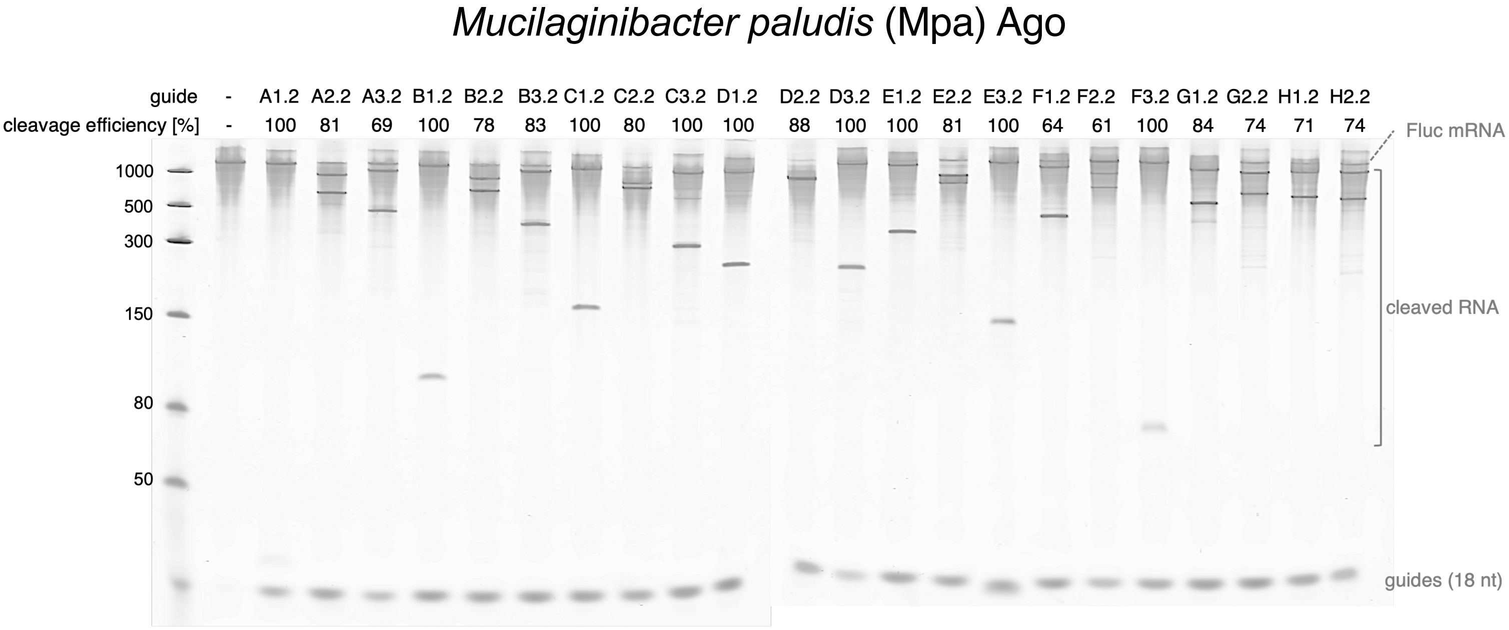

Supplemental Figure 4

A

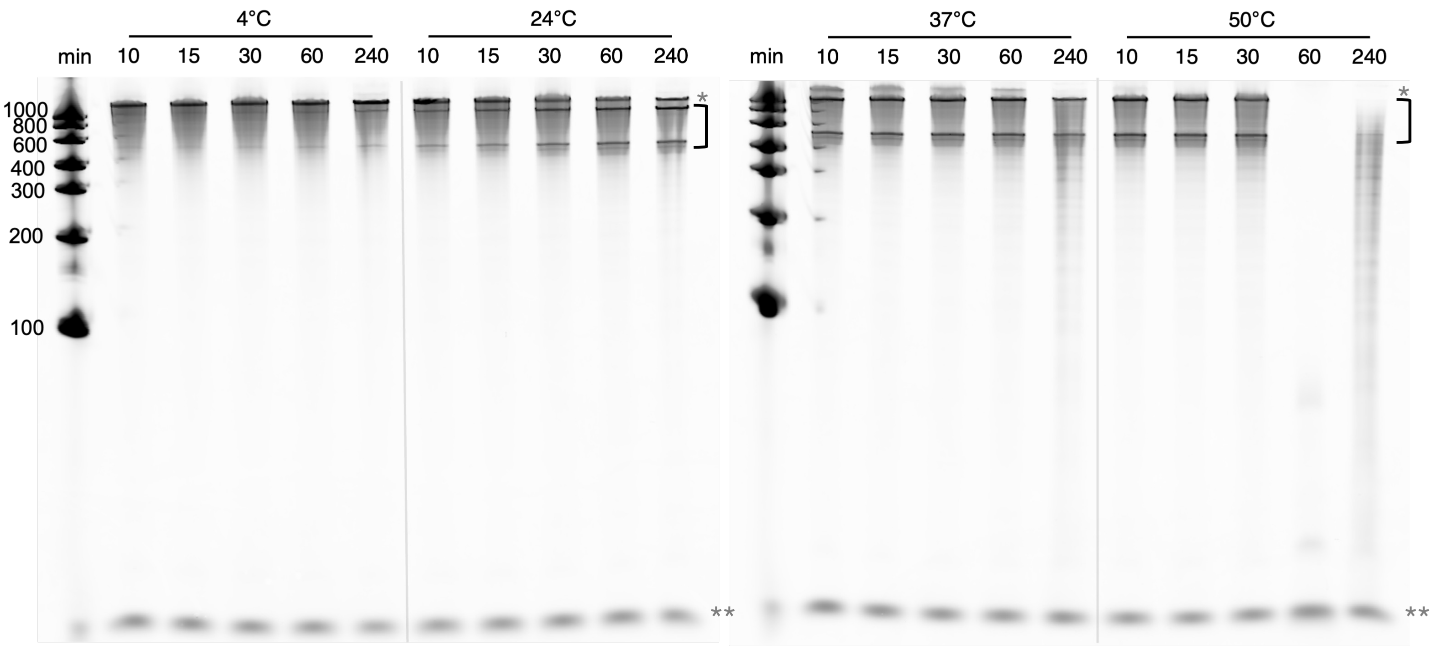


B

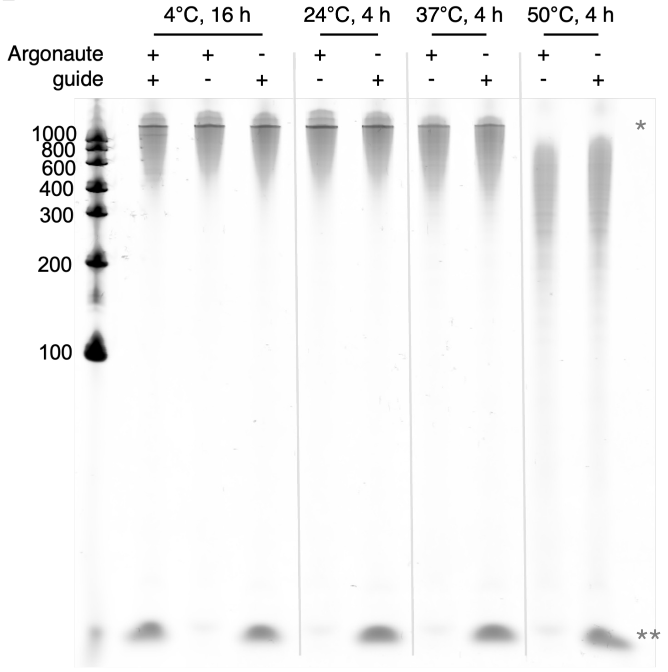

Supplemental Figure 5

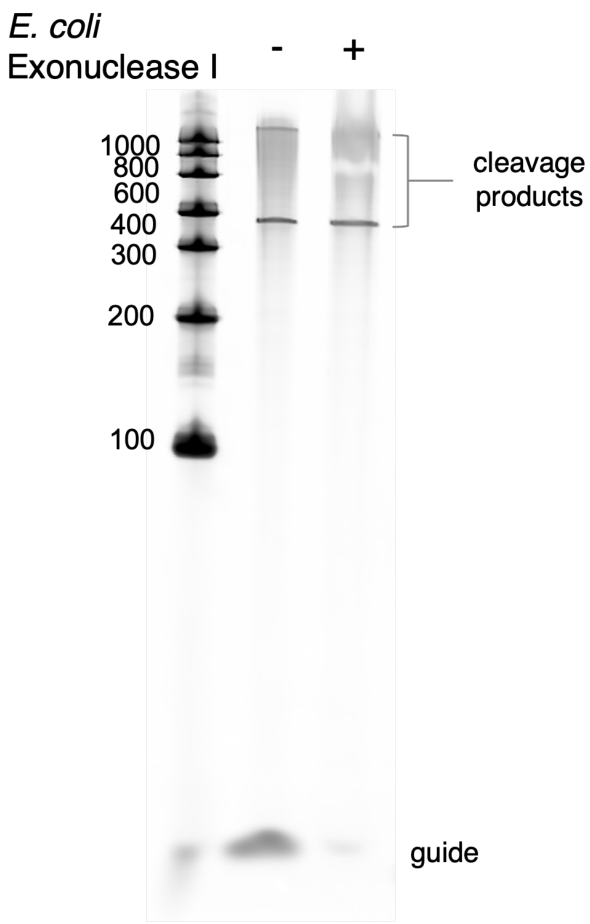

Supplemental Figure 6

### Sequence coverage of uniquely mappable fragments

#### Experiment 2: 79.2% Sequence Coverage

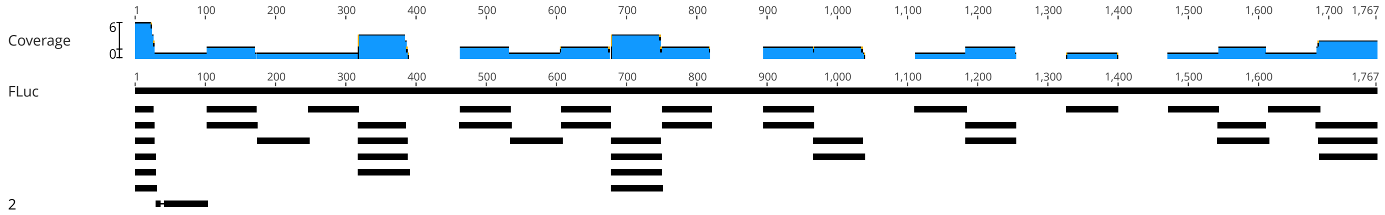


#### Experiment 3: 83.5% Sequence Coverage

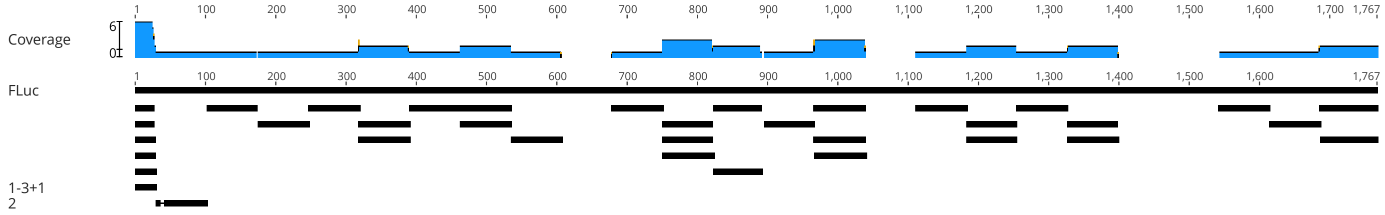


#### Experiment 4: 82.9% Sequence Coverage

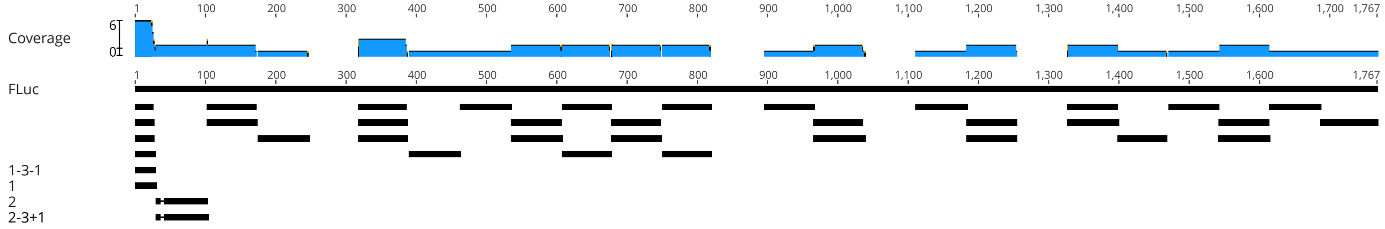


#### Experiment 5: 87.7% Sequence Coverage

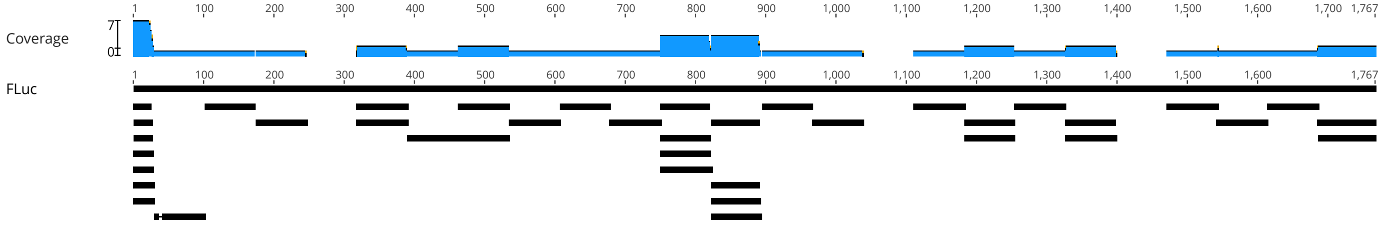

Supplemental Figure 7

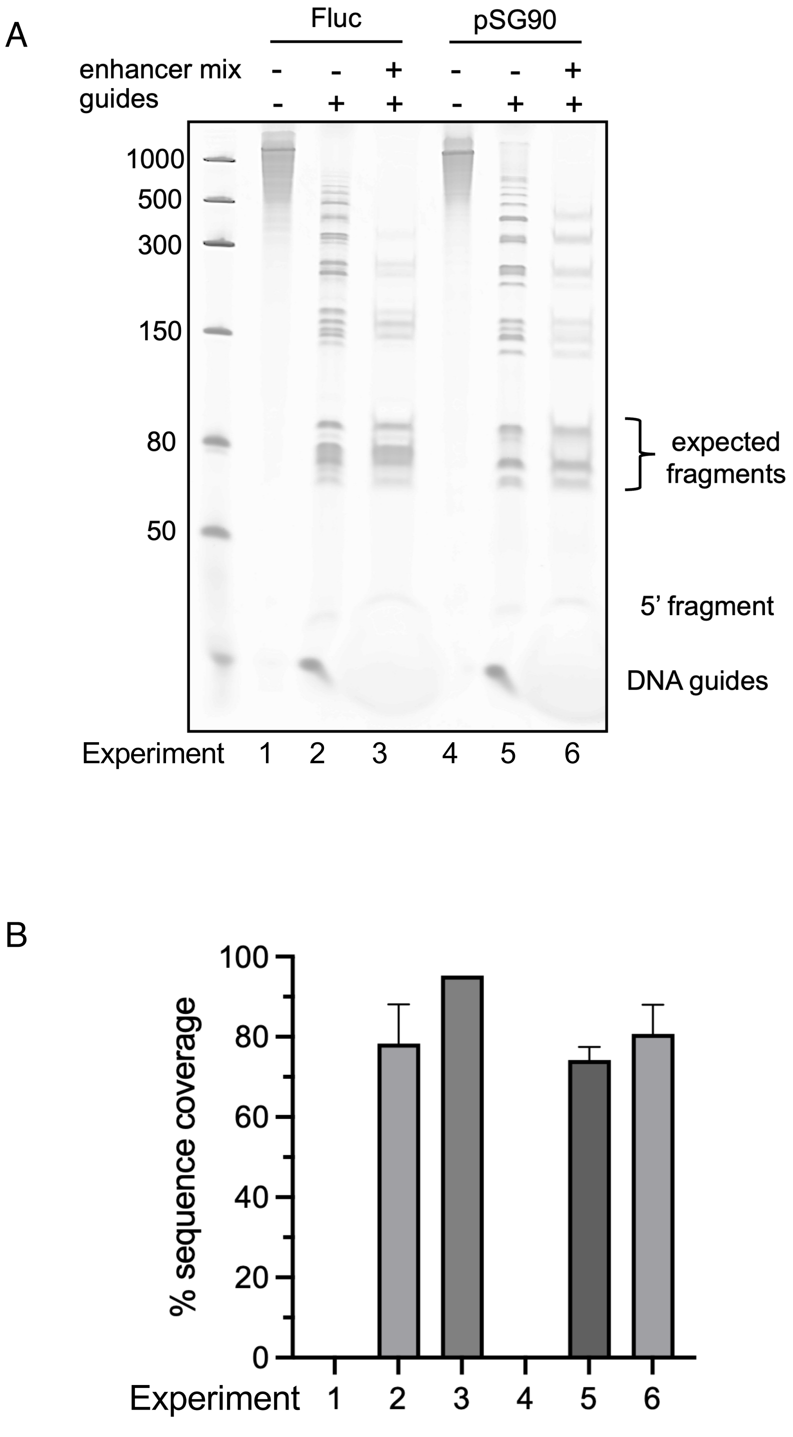

Supplemental Figure 7

C Sequence coverage of uniquely mappable fragments

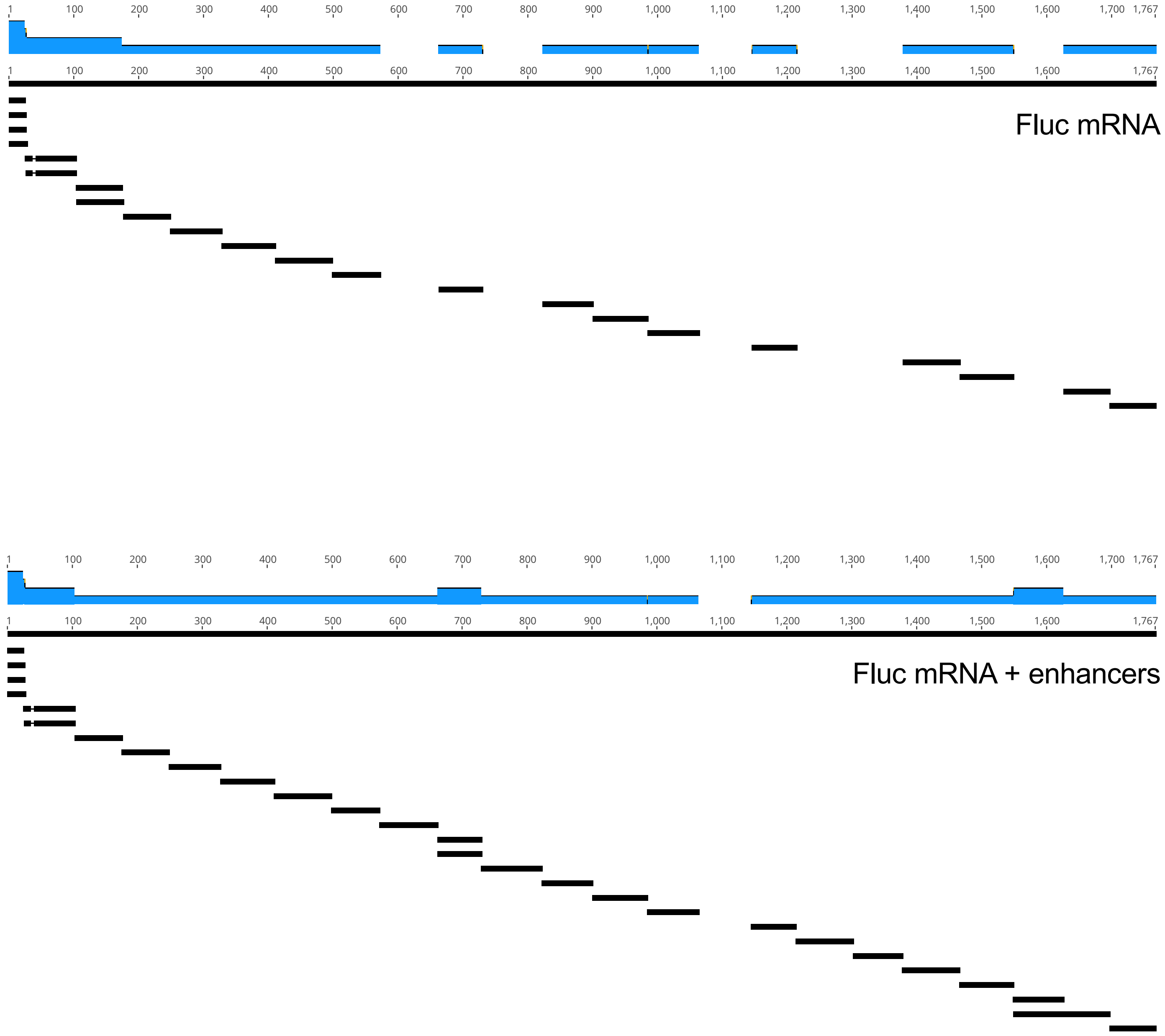

Supplemental Figure 7

D Sequence coverage of uniquely mappable fragments

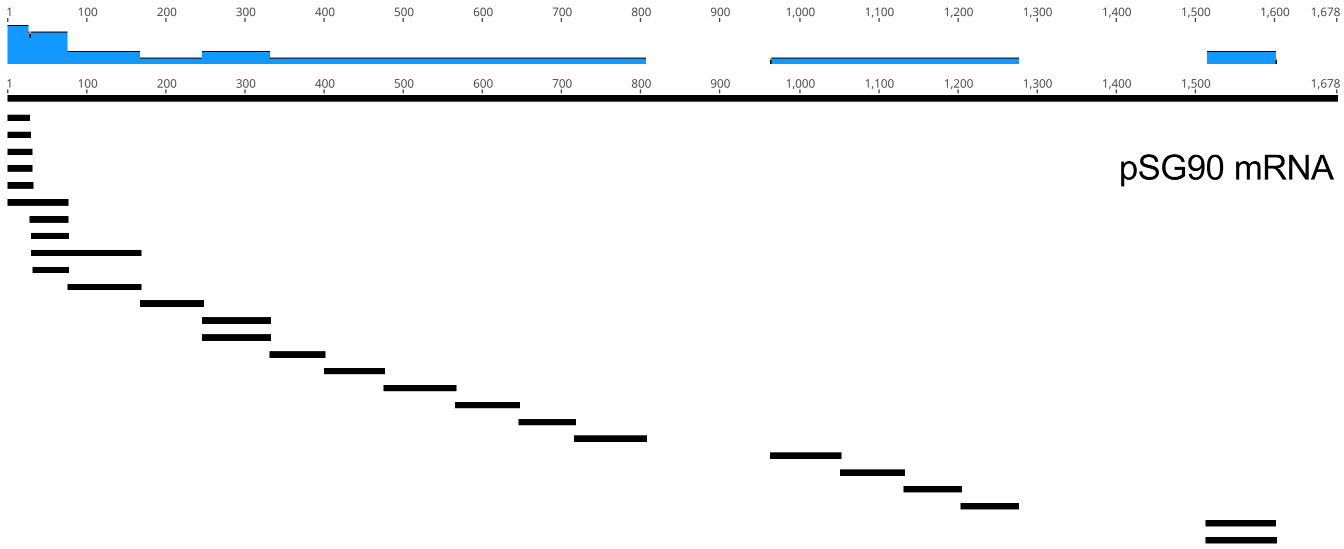


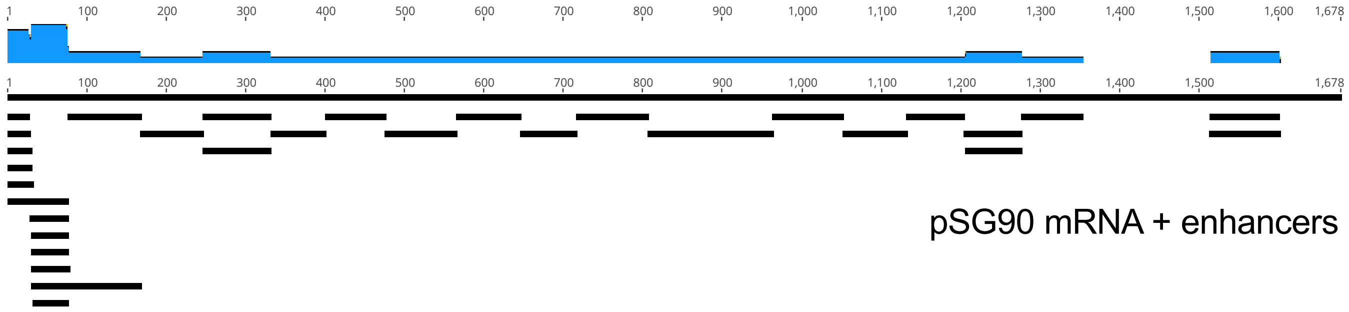

Supplemental Figure 8

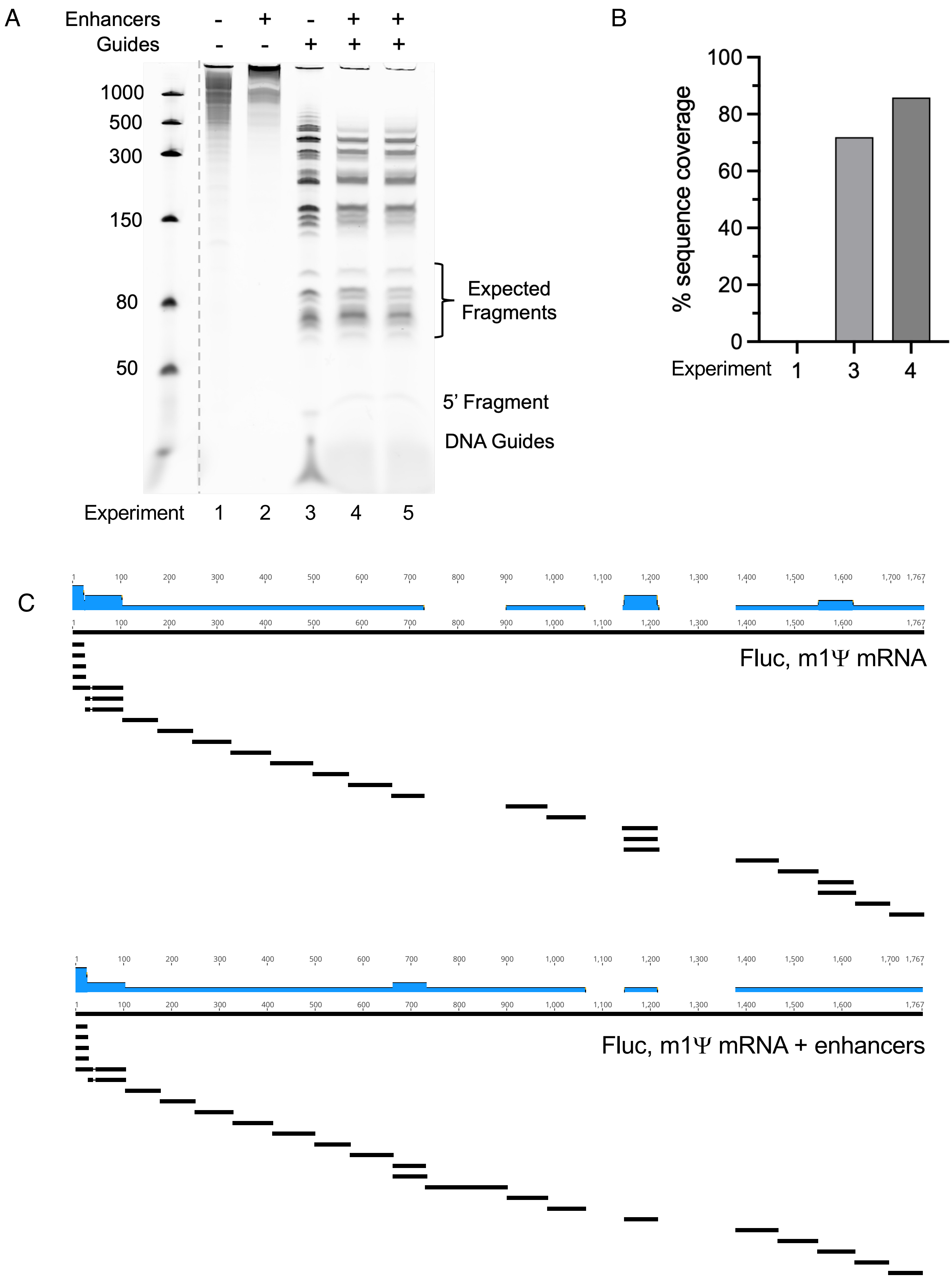

### Supplemental Figure 9

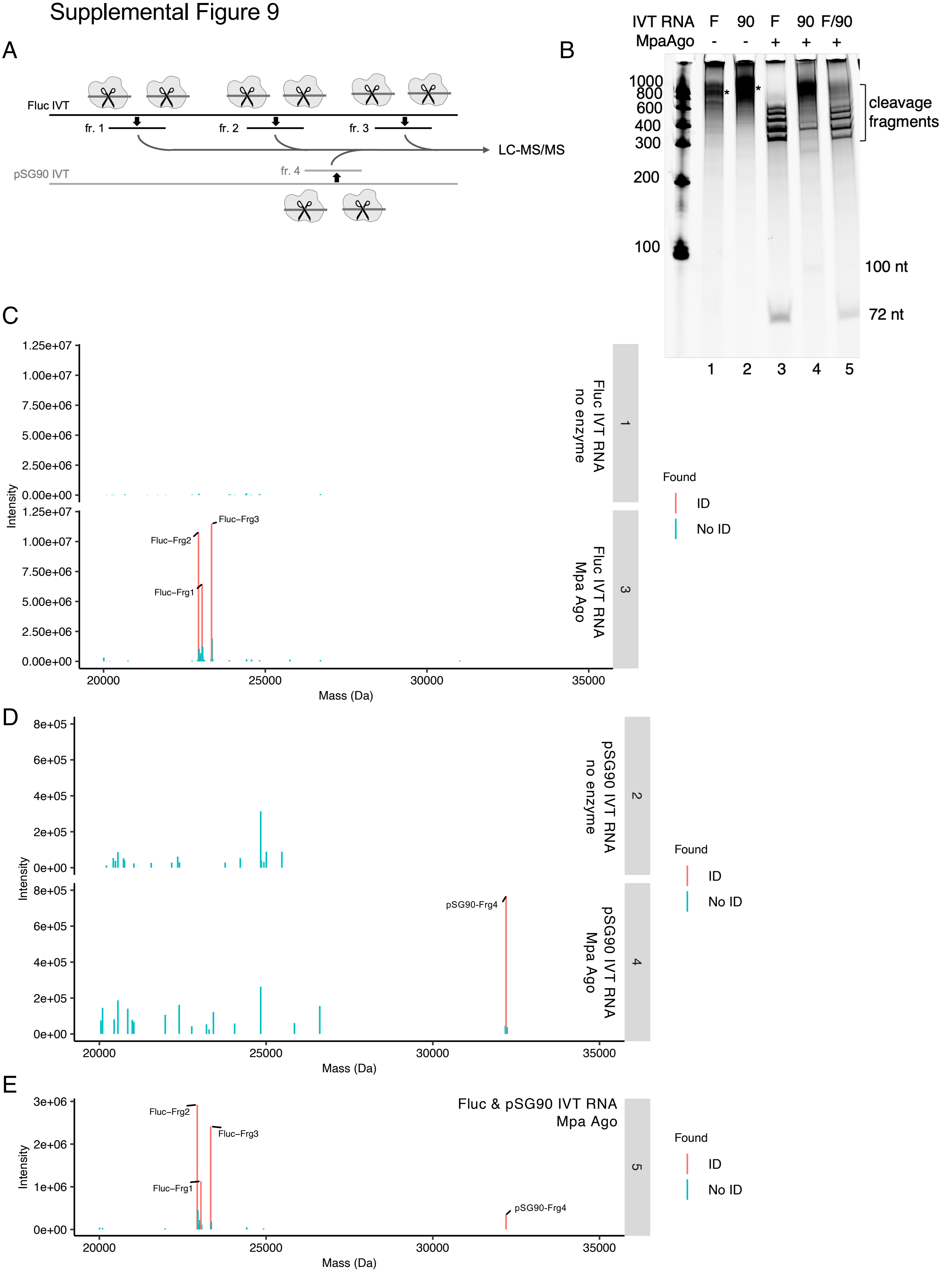

Supplemental Figure 10

A

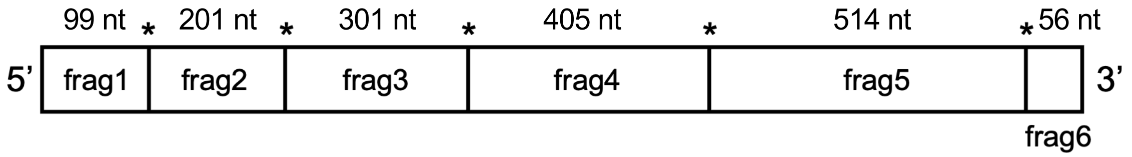


B

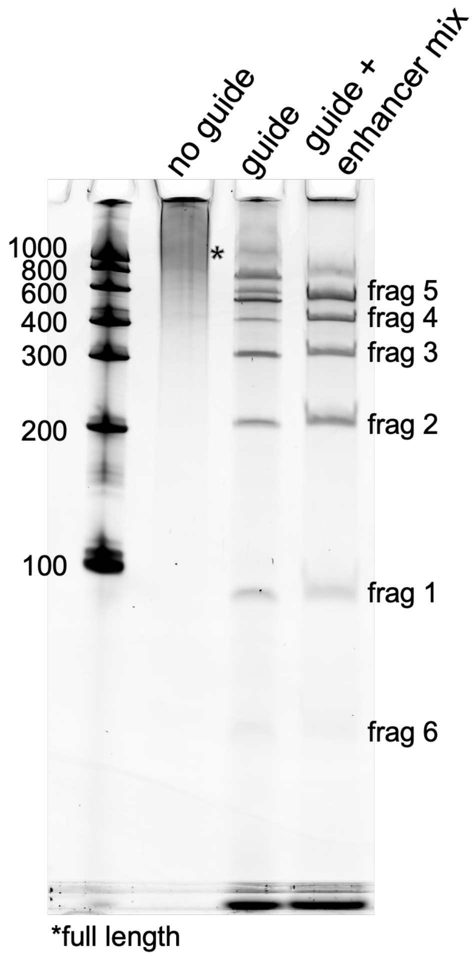

Supplemental Figure 11

A

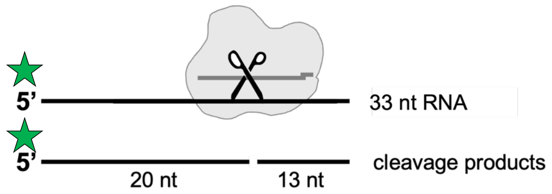


B

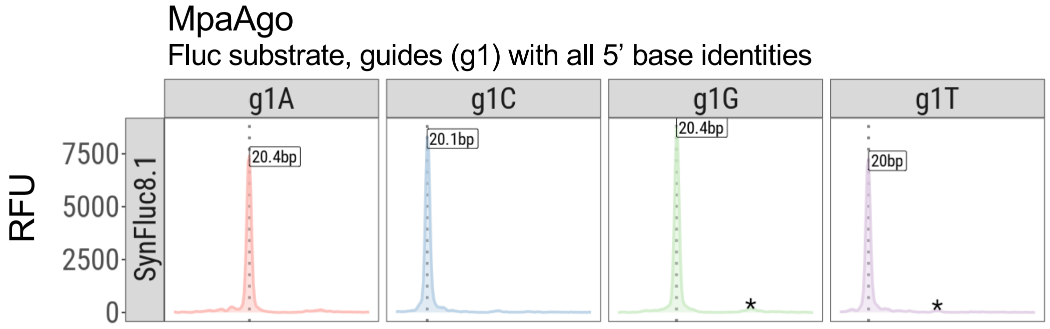

Supplemental Figure 12

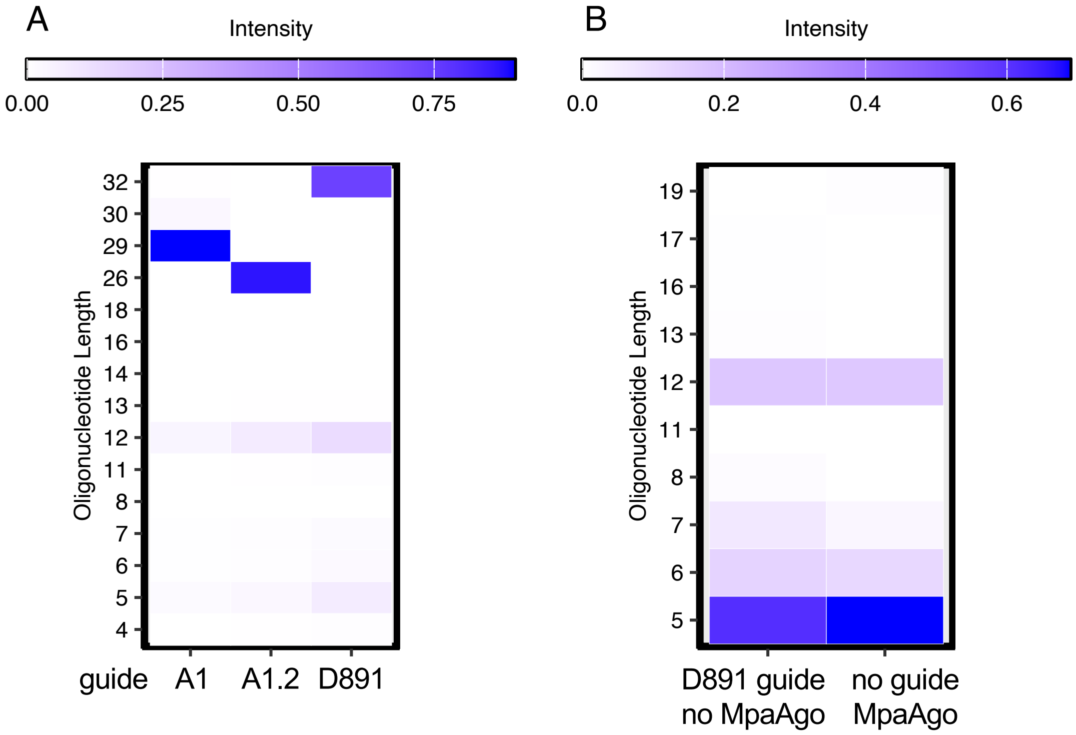


**Fluc mRNA**
GGGUCUAGAAAUAAUUUUGUUUAACUUUAAGAAGGAGA… (wt)
GGGUCUAGAAAUAAUUUUGUUUAACUUUA| (A1 guide; 29mer)
GGGUCUAGAAAUAAUUUUGUUUAACU| (A1.2 guide; 26mer)
GGGUCUAGAAAUAAUUUUGUUUAACUUUAAGA| (D891 guide; 32mer)

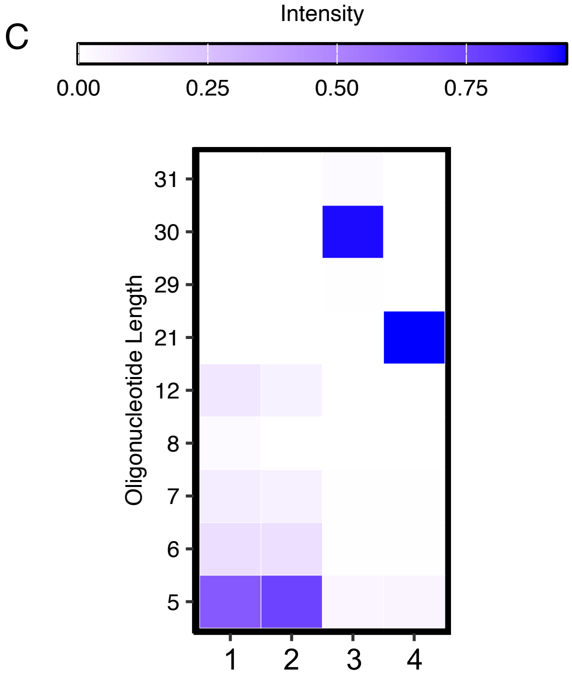


1. D831 guide, no MpaAgo
2. no guide, MpaAgo
3. D831 guide, MpaAgo
4. D892 guide, MpaAgo

**pSG90 mRNA**
GGUGACGGCGUAGUACACACUAUUGAAUCAAACAGCCGACCAAUU… (wt)
GGUGACGGCGUAGUACACACUAUUGAAUCA| (30mer)
GGUGACGGCGUAGUACACACU| (21mer)

Supplemental Figure 13

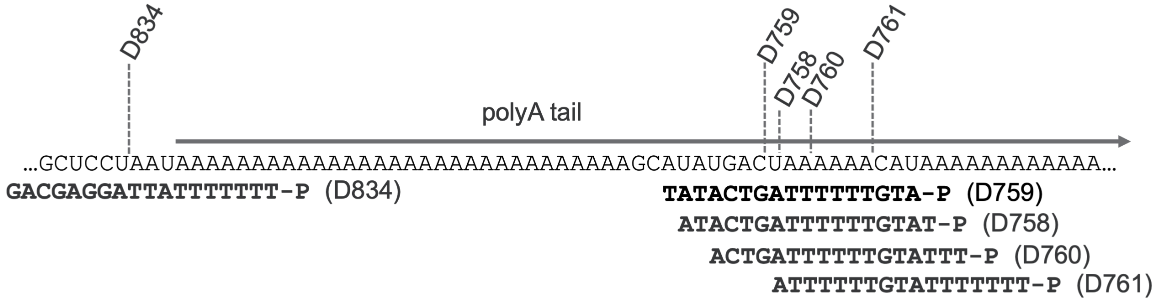

Supplemental Figure 14

A

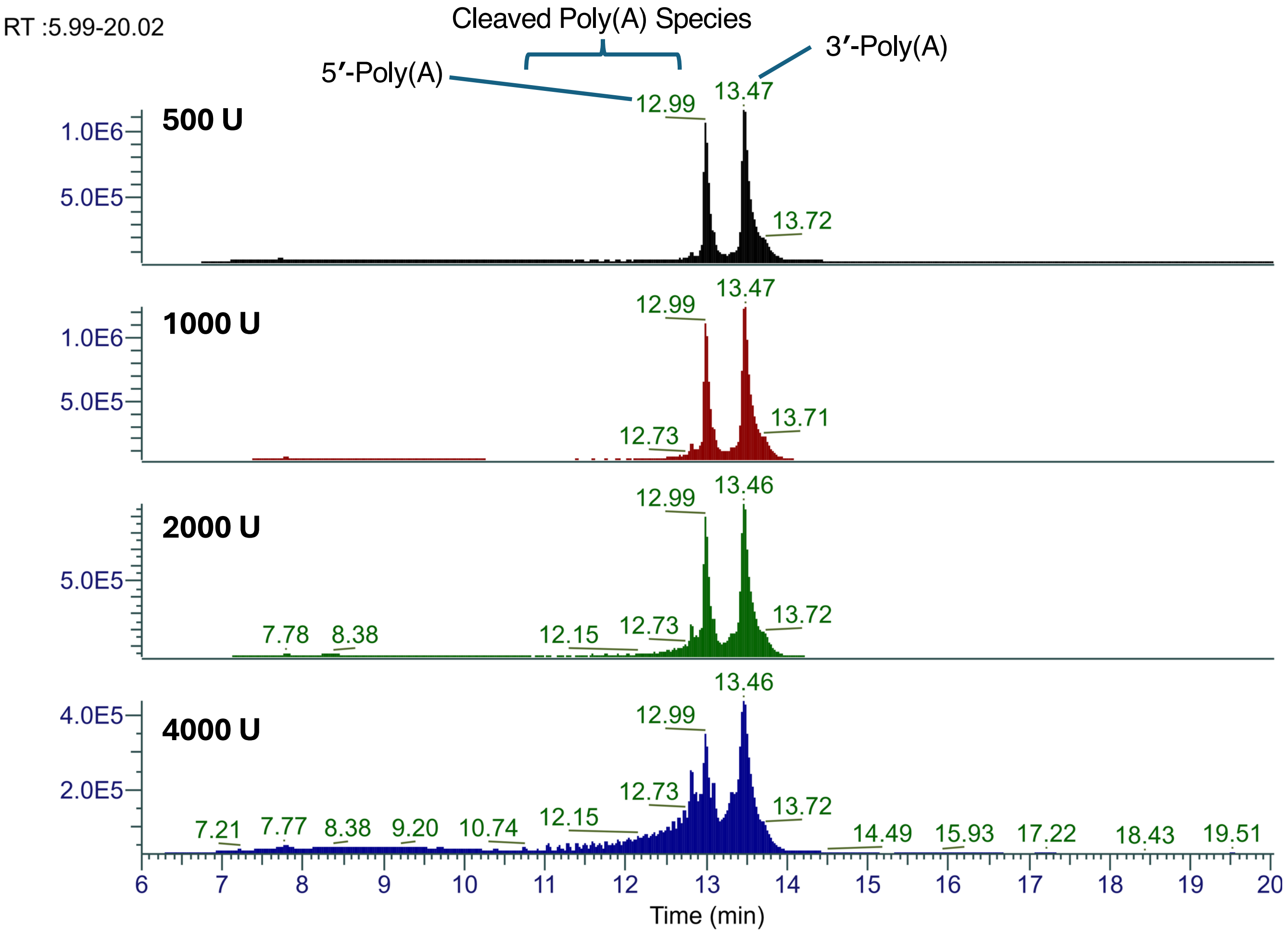


B

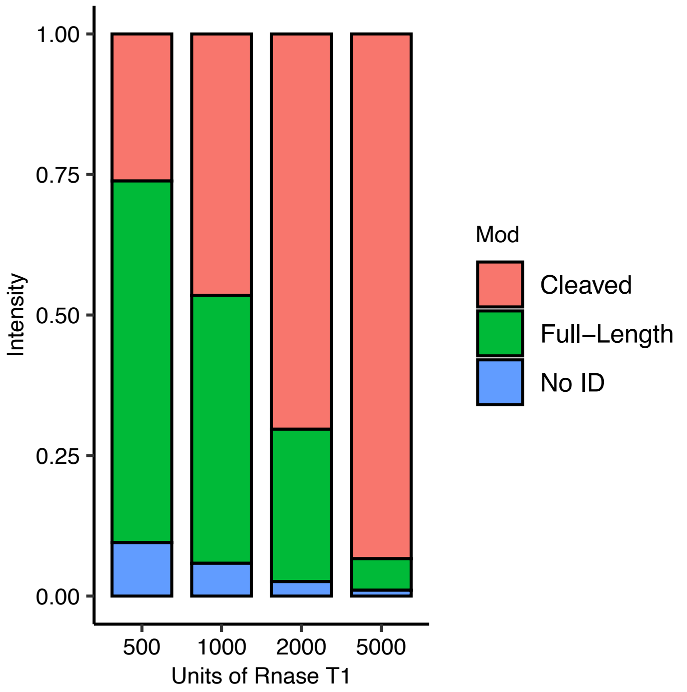

### Supplemental Table 1

<table>
<tr><th>Experimental Mass (Da)</th><th>Theoretical Mass (Da)</th><th>Intensity</th><th>Sequences</th></tr>
<tr><td>5257.864</td><td>5257.8527</td><td>3.30E+06</td><td>{dA}{dT}{dA}{dC}{dT}{dG}{dA}{dT}{dT}{dT}{dT}{dT}{dT}{dT}{dG}{dT}{dA}{dT}|-p|</td></tr>
<tr><td>17975.779</td><td>17975.7292</td><td>1.23E+07</td><td>rArArArArArArCrArUrArArArArArArArArArArArArArArArArArArArArArArArArArArArArArArArArArArArArArGrCrUrArUrCrGrGrArUrC|-p||Minus-A|</td></tr>
<tr><td>18280.799</td><td>18280.7705</td><td>1.01E+07</td><td>rArArArArArArCrArUrArArArArArArArArArArArArArArArArArArArArArArArArArArArArArArArArArArArArArGrCrUrArUrCrGrGrArUrCrC|-p||Minus-A|</td></tr>
<tr><td>18304.787</td><td>18304.7817</td><td>7.57E+07</td><td>rArArArArArArCrArUrArArArArArArArArArArArArArArArArArArArArArArArArArArArArArArArArArArArArArGrCrUrArUrCrGrGrArUrC|-p|</td></tr>
<tr><td>18609.848</td><td>18609.823</td><td>7.75E+07</td><td>rArArArArArArCrArUrArArArArArArArArArArArArArArArArArArArArArArArArArArArArArArArArArArArArArGrCrUrArUrCrGrGrArUrCrC|-p|</td></tr>
<tr><td>18633.818</td><td>18633.8342</td><td>6.49E+07</td><td>rArArArArArArCrArUrArArArArArArArArArArArArArArArArArArArArArArArArArArArArArArArArArArArArArGrCrUrArUrCrGrGrArUrC|-p||Plus-A|</td></tr>
<tr><td>18938.853</td><td>18938.8755</td><td>5.42E+07</td><td>rArArArArArArCrArUrArArArArArArArArArArArArArArArArArArArArArArArArArArArArArArArArArArArArArGrCrUrArUrCrGrGrArUrCrC|-p||Plus-A|</td></tr>
<tr><td>18962.924</td><td>18962.8867</td><td>2.57E+07</td><td>rArArArArArArCrArUrArArArArArArArArArArArArArArArArArArArArArArArArArArArArArArArArArArArArArGrCrUrArUrCrGrGrArUrC|-p||Plus-2A|</td></tr>
<tr><td>19267.943</td><td>19267.928</td><td>1.21E+07</td><td>rArArArArArArCrArUrArArArArArArArArArArArArArArArArArArArArArArArArArArArArArArArArArArArArArGrCrUrArUrCrGrGrArUrCrC|-p||Plus-2A|</td></tr>
<tr><td>19291.94</td><td>19291.9392</td><td>8.35E+06</td><td>rArArArArArArCrArUrArArArArArArArArArArArArArArArArArArArArArArArArArArArArArArArArArArArArArGrCrUrArUrCrGrGrArUrC|-p||Plus-3A|</td></tr>
<tr><td>19597.004</td><td>19596.9805</td><td>5.48E+06</td><td>rArArArArArArCrArUrArArArArArArArArArArArArArArArArArArArArArArArArArArArArArArArArArArArArArGrCrUrArUrCrGrGrArUrCrC|-p||Plus-3A|</td></tr>
</table>

### Supplemental Table 2

<table>
<tr><th>Experimental Mass (Da)</th><th>Theoretical Mass (Da)</th><th>Intensity</th><th>Sequences</th></tr>
<tr><td>30677.61</td><td>30677.6081</td><td>2.86E+07</td><td>rUrArArUrArArArArArArArArArArArArArArArArArArArArArArArGrCrArUrArUrGrArCrUrArArArArArArCrArUrArArArArArArArArArArArArArArArArArArArArArArArArArArArArArArArArArArArArArGrCrUrArUrCrGrGrArUrC|-p||Plus-2A|</td></tr>
<tr><td>30676.594</td><td>30676.6241</td><td>1.46E+07</td><td>rArArUrArArArArArArArArArArArArArArArArArArArArArArArGrCrArUrArUrGrArCrUrArArArArArArCrArUrArArArArArArArArArArArArArArArArArArArArArArArArArArArArArArArArArArArArArGrCrUrArUrCrGrGrArUrCrC|-p||Plus-2A|</td></tr>
<tr><td>30371.587</td><td>30371.5828</td><td>3.05E+07</td><td>rArArUrArArArArArArArArArArArArArArArArArArArArArArArGrCrArUrArUrGrArCrUrArArArArArArCrArUrArArArArArArArArArArArArArArArArArArArArArArArArArArArArArArArArArArArArGrCrUrArUrCrGrGrArUrC|-p||Plus-2A|</td></tr>
<tr><td>30348.569</td><td>30348.5556</td><td>3.37E+06</td><td>rUrArArUrArArArArArArArArArArArArArArArArArArArArArArArGrCrArUrArUrGrArCrUrArArArArArArCrArUrArArArArArArArArArArArArArArArArArArArArArArArArArArArArArArArArArArArArGrCrUrArUrCrGrGrArUrC|-p||Plus-A|</td></tr>
<tr><td>30347.617</td><td>30347.5716</td><td>8.69E+07</td><td>rArArUrArArArArArArArArArArArArArArArArArArArArArArArGrCrArUrArUrGrArCrUrArArArArArArCrArUrArArArArArArArArArArArArArArArArArArArArArArArArArArArArArArArArArArArArGrCrUrArUrCrGrGrArUrCrC|-p||Plus-A|</td></tr>
<tr><td>30042.505</td><td>30042.5303</td><td>6.35E+07</td><td>rArArUrArArArArArArArArArArArArArArArArArArArArArArArGrCrArUrArUrGrArCrUrArArArArArArCrArUrArArArArArArArArArArArArArArArArArArArArArArArArArArArArArArArArArArArArGrCrUrArUrCrGrGrArUrC|-p||Plus-A|</td></tr>
<tr><td>30019.526</td><td>30019.5031</td><td>8.29E+07</td><td>rUrArArUrArArArArArArArArArArArArArArArArArArArArArArArGrCrArUrArUrGrArCrUrArArArArArArCrArUrArArArArArArArArArArArArArArArArArArArArArArArArArArArArArArArArArArArArGrCrUrArUrCrGrGrArUrC|-p|</td></tr>
<tr><td>30018.538</td><td>30018.5191</td><td>1.94E+07</td><td>rArArUrArArArArArArArArArArArArArArArArArArArArArArArGrCrArUrArUrGrArCrUrArArArArArArCrArUrArArArArArArArArArArArArArArArArArArArArArArArArArArArArArArArArArArArArGrCrUrArUrCrGrGrArUrCrC|-p|</td></tr>
<tr><td>29713.485</td><td>29713.4778</td><td>7.31E+07</td><td>rArArUrArArArArArArArArArArArArArArArArArArArArArArArGrCrArUrArUrGrArCrUrArArArArArArCrArUrArArArArArArArArArArArArArArArArArArArArArArArArArArArArArArArArArArArArGrCrUrArUrCrGrGrArUrC|-p|</td></tr>
<tr><td>5611.924</td><td>5611.91167</td><td>2.92E+06</td><td>{dG}{dA}{dC}{dG}{dA}{dG}{dG}{dA}{dT}{dT}{dA}{dT}{dT}{dT}{dT}{dT}{dT}{dT}|-p|</td></tr>
<tr><td>5531.957</td><td>5531.94537</td><td>1.85E+06</td><td>{dG}{dA}{dC}{dG}{dA}{dG}{dG}{dA}{dT}{dT}{dA}{dT}{dT}{dT}{dT}{dT}{dT}{dT}</td></tr>
</table>

Supplemental Table 3

| | |
|---|---|
| Fluc mRNA | GGGUCUAGAAAUAAUUUUGUUUAACUUUAAGAAGGAGAUAUAACCAUGAAAAUCGAAGAAGGUAAAGGUCACCAUCACCAUCACCACGGAUCCAUGGAAGACGCCAAAAACAUAAAGAAAGGCCCGGCGCCAUUCUAUCCUCUAGAGGAUGGAACCGCUGGAGAGCAACUGCAUAAGGCUAUGAAGAGAUACGCCCUGGUUCCUGGAACAAUUGCUUUUACAGAUGCACAUAUCGAGGUGAACAUCACGUACGCGGAAUACUUCGAAAUGUCCGUUCGGUUGGCAGAAGCUAUGAAACGAUAUGGGCUGAAUACAAAUCACAGAAUCGUCGUAUGCAGUGAAAACUCUCUUCAAUUCUUUAUGCCGGUGUUGGGCGCGUUAUUUAUCGGAGUUGCAGUUGCGCCCGCGAACGACAUUUAUAAUGAACGUGAAUUGCUCAACAGUAUGAACAUUUCGCAGCCUACCGUAGUGUUUGUUUCCAAAAAGGGGUUGCAAAAAAUUUUGAACGUGCAAAAAAAAUUACCAAUAAUCCAGAAAAUUAUUAUCAUGGAUUCUAAAACGGAUUACCAGGGAUUUCAGUCGAUGUACACGUUCGUCACAUCUCAUCUACCUCCCGGUUUUAAUGAAUACGAUUUUGUACCAGAGUCCUUUGAUCGUGACAAAACAAUUGCACUGAUAAUGAAUUCCUCUGGAUCUACUGGGUUACCUAAGGGUGUGGCCCUUCCGCAUAGAACUGCCUGCGUCAGAUUCUCGCAUGCCAGAGAUCCUAUUUUUGGCAAUCAAAUCAUUCCGGAUACUGCGAUUUUAAGUGUUGUUCCAUUCCAUCACGGUUUUGGAAUGUUUACUACACUCGGAUAUUUGAUAUGUGGAUUUCGAGUCGUCUUAAUGUAUAGAUUUGAAGAAGAGCUGUUUUUACGAUCCCUUCAGGAUUACAAAAUUCAAAGUGCGUUGCUAGUACCAACCCUAUUUUCAUUCUUCGCCAAAAGCACUCUGAUUGACAAAUACGAUUUAUCUAAUUUACACGAAAUUGCUUCUGGGGGCGCACCUCUUUCGAAAGAAGUCGGGGAAGCGGUUGCAAAACGCUUCCAUCUUCCAGGGAUACGACAAGGAUAUGGGCUCACUGAGACUACAUCAGCUAUUCUGAUUACACCCGAGGGGGAUGAUAAACCGGGCGCGGUCGGUAAAGUUGUUCCAUUUUUUGAAGCGAAGGUUGUGGAUCUGGAUACCGGGAAAACGCUGGGCGUUAAUCAGAGAGGCGAAUUAUGUGUCAGAGGACCUAUGAUUAUGUCCGGUUAUGUAAACAAUCCGGAAGCGACCAACGCCUUGAUUGACAAGGAUGGAUGGCUACAUUCUGGAGACAUAGCUUACUGGGACGAAGACGAACACUUCUUCAUAGUUGACCGCUUGAAGUCUUUAAUUAAAUACAAAGGAUAUCAGGUGGCCCCCGCUGAAUUGGAAUCGAUAUUGUUACAACACCCCAACAUCUUCGACGCGGGCGUGGCAGGUCUUCCCGACGAUGACGCCGGUGAACUUCCCGCCGCCGUUGUUGUUUUGGAGCACGGAAAGACGAUGACGGAAAAAGAGAUCGUGGAUUACGUCGCCAGUCAAGUAACAACCGCGAAAAAGUUGCGCGGAGGAGUUGUGUUUGUGGACGAAGUACCGAAAGGUCUUACCGGAAAACUCGACGCAAGAAAAAUCAGAGAGAUCCUCAUAAAGGCCAAGAAGGGCGGAAAGUCCAAACUCGAGUAAGGUUAACCUGCAGGAGG |
| pSG90 mRNA | GGUGACGGCGUAGUACACACUAUUGAAUCAAACAGCCGACCAAUUGCACUACCAUCACAAUGGAGAAGCCAGUAGUAAACGUAGACGUAGACCCCCAGAGUCCGUUUGUCGUGCAACUGCAAAAAAGCUUCCCGCAAUUUGAGGUAGUAGCACAGCAGGUCACUCCAAAUGACCAUGCUAAUGCCAGAGCAUUUUCGCAUCUGGCCAGUAAACUAAUCGAGCUGGAGGUUCCUACCACAGCGACGAUCUUGGACAUAGGCAGCGCACCGGCUCGUAGAAUGUUUUCCGAGCACCAGUAUCAUUGUGUCUGCCCCAUGCGUAGUCCAGAAGACCCGGACCGCAUGAUGAAAUACGCCAGUAAACUGGCGGAAAAAGCGUGCAAGAUUACAAACAAGAACUUGCAUGAGAAGAUUAAGGAUCUCCGGACCGUACUUGAUACGCCGGAUGCUGAAACACCAUCGCUCUGCUUUCACAACGAUGUUACCUGCAACAUGCGUGCCGAAUAUUCCGUCAUGCAGGACGUGUAUAUCAACGCUCCCGGAACUAUCUAUCAUCAGGCUAUGAAAGGCGUGCGGACCCUGUACUGGAUUGGCUUCGACACCACCCAGUUCAUGUUCUCGGCUAUGGCAGGUUCGUACCCUGCGUACAACACCAACUGGGCCGACGAGAAAGUCCUUGAAGCGCGUAACAUCGGACUUUGCAGCACAAAGCUGAGUGAAGGUAGGACAGGAAAAUUGUCGAUAAUGAGGAAGAAGGAGUUGAAGCCCGGGUCGCGGGUUUAUUUCUCCGUAGGAUCGACACUUUAUCCAGAACACAGAGCCAGCUUGCAGAGCUGGCAUCUUCCAUCGGUGUUCCACUUGAAUGGAAAGCAGUCGUACACUUGCCGCUGUGAUACAGUGGUGAGUUGCGAAGGCUACGUAGUGAAGAAAAUCACCAUCAGUCCCGGGAUCACGGGAGAAACCGUGGGAUACGCGGUUACACACAAUAGCGAGGGCUUCUUGCUAUGCAAAGUUACUGACACAGUAAAAGGAGAACGGGUAUCGUUCCCUGUGUGCACGUACAUCCCGGCCACCAUAUGCGAUCAGAUGACUGGUAUAAUGGCCACGGAUAUAUCACCUGACGAUGCACAAAAACUUCUGGUUGGGCUCAACCAGCGAAUUGUCAUUAACGGUAGGACUAACAGGAACACCAACACCAUGCAAAAUUACCUUCUGCCGAUCAUAGCACAAGGGUUCAGCAAAUGGGCUAAGGAGCGCAAGGAUGAUCUUGAUAACGAGAAAAUGCUGGGUACUAGAGAACGCAAGCUUACGUAUGGCUGCUUGUGGGCGUUUCGCACUAAGAAAGUACAUUCGUUUUAUCGCCCACCUGGAACGCAGACCUGCGUAAAAGUCCCAGCCUCUUUUAGCGCUUUUCCCAUGUCGUCCGUAUGGACGACCUCUUUGCCCAUGUCGCUGAGGCAGAAAUUGAAACUGGCAUUGCAACCAAAGAAGGAGGAAAAACUGCUGCAGGUCUCGGAGGAAUUAGUCAUGGAGGCCAUUUUCCGAAUCGGAUUUUGUUUUAAUAUUUCAAAAAAAAAAAAAAAAAAAAAAAAAAAAAGCAUAUGACUAAAAAAAAAAAAAAAAAAAAAAAAAAAAAAAAAAAAAAAAAAAAAAAAAAAAAAAAAAAAAAAAAAAA |
| pSG95 mRNA | GGGCUUGCUUGUUCUUUUUGCAGAAGCUCAGAAUAAACGCUCAACUUUGGCACCGCCGCACAACCGGAUGCGCCGAGCACUGCAGCCUCAACGAGAACAUCACCGUACCGGACACAAAGGUCAACUUUACGCAUGGAAGAGAAUGGAAGUAGGACAGCAGGCCGUCGAAGUGUGCAGGGGCUCGCGCUGCUGAGCGAGGCGGUGCUGCGGGGCCAGGCCCUCCUCGUCAACAGCAGCCAGCCGUGGGAGCCCCUCCAACUGCACGUCGACAAAGCGGUGAGCGGGCUCCGCAGCCUGACGACGCUGCUGCGGGCCCUGGGCGCACAAAAGGAGGCCAUCAGCCCGCCCGACGCGGCCAGCGCGGCACCCCCCGAACGAUCACCGCGGACACGUUCAGGAAGCUGUUAGAGUGUACAGCAACUUCCUCGCGGAAAGCUGAAACUGUACACCGGCGAAGCGUGCAGGACAGGGGACCGCUAGGACUGACUAGGAUUGGUUACCACUAAACCAGCCUCAAGAACACCCGAAUGGAGUCUCUAAGCUACAUAAUACCAACUUACACUUACAACAUGUGUCCCCCAAACUGUAGCCAUUCGUAUCUGCUCCUAAUAAAAAAAAAAAAAAAAAAAAAAAAAAAAGCAUAUGACUAAAAAACAUAAAAAAAAAAAAAAAAAAAAAAAAAAAAAAAAAAAAAAAAAAAAAAAAAAAAAAAAAAAAGCU |

| | |
|---|---|
| pSG120 mRNA | GGGCUUGCUUGUUCUUUUUGCAGAAGCUCAGAAUAAACGCUCAACUUUGGCACCGCCGCACAACCGGAUGC<br>GCCGAGCACUGCAGCCUCAACGAGAACAUCACCGUACCGGACACAAAGGUCAACUUUACGCAUGGAAGAGAA<br>UGGAAGUAGGACAGCAGGCCGUCGAAGUGUGCAGGGGCUCGCGCUGCUGAGCGAGGCGGUGCUGCGGGG<br>CCAGGCCCUCCUCGUCAACAGCAGCCAGCCGUGGGAGCCCCUCCAACUGCACGUCGACAAAGCGGUGAGCG<br>GGCUCCGCAGCCUGACGACGCUGCUGCGGGCCCUGGGCGCACAAAAGGAGGCCAUCAGCCCGCCCGACGC<br>GGCCAGCGCGGCACCCCCCGAACGAUCACCGCGGACACGUUCAGGAAGCUGUUAGAGUGUACAGCAACUUC<br>CUCGCGGAAAGCUGAAACUGUACACCGGCGAAGCGUGCAGGACAGGGGACCGCUAGGACUGACUAGGAUUG<br>GUUACCACUAAACCAGCCUCAAGAACACCCGAAUGGAGUCUCUAAGCUACAUAAUACCAACUUACACUUACAA<br>CAUGUGUCCCCCAAACUGUAGCCAUUCGUAUCUGCUCCUAAUAAAAAAAAAAAAAAAAAAAAAGCAUAUGAC<br>UAAAAAACAUAAAAAAAAAAAAAAAAAAAAAAAAAAAAAAAAAAAAAGCUAUCGGAUC |
| pSG106 mRNA | GGUGACGGCGUAGUACACACUAUUGAAUCAAACAGCCGACCAAUUGCACUACCAUCACAAUGGAGAAGCCAG<br>UAGUAAACGUAGACGUAGACCUGUCAUUAACGGUAGGCGUGCAACUGCAAAAAAGCUUCCCGCAAUUUGAGG<br>UAGUAGCACAGCAGGUCACUCCAAAUGACCAUGCUAAUGCCAGAGCAUUUUCGCAUCUGGCCAGUAAACUAA<br>UCGAGCUGGAGGUUCCUACCACAGCGACGAUCUUGGACAUAGGCAGCGCACCGGCUCGUAGAAUGUUUUCC<br>GAGGACCUGUCAUUAACGGUAGGCCCCAUGCGUAGUCCAGAAGACCCGGACCGCAUGAUGAAAUACGCCAG<br>UAAACUGGCGGAAAAAGCGUGCAAGAUUACAAACAAGAACUUGCAUGAGAAGAUUAAGGAUCUCCGGACCGU<br>ACUUGAUACGCCGGAUGCUGAAACACCAUCGCUCUGCUUUCACAACGAUGUUACCUGCAACAUGCGUGCCG<br>AAUAUUCCGUCAUGCAGGACGUGUAUAUCAACGCUCCCGGAACUAUCUAUCAUCAGGCUAUGAAAGGCGUGC<br>GGACCCUGUACUGGAUUGGACCUGUCAUUAACGGUAGGGUUCAUGUUCUCGGCUAUGGCAGGUUCGUACC<br>CUGCGUACAACACCAACUGGGCCGACGAGAAAGUCCUUGAAGCGCGUAACAUCGGACUUUGCAGCACAAAGC<br>UGAGUGAAGGUAGGACAGGAAAAUUGUCGAUAAUGAGGAAGAAGGAGUUGAAGCCCGGGUCGCGGGUUUAU<br>UUCUCCGUAGGAUCGACACUUUAUCCAGAACACAGAGCCAGCUUGCAGAGCUGGCAUCUUCCAUCGGUGUU<br>CCACUUGAAUGGAAAGCAGUCGUACACUUGCCGCUGUGAUACAGUGGUGAGUUGCGAAGGCUACGUAGUGA<br>AGAAAAUCACCAUCAGUCCCGGGAUCACGGGAGAAACCGUGGGAUACGCGGUUACACACAAUAGCGAGGACC<br>UGUCAUUAACGGUAGGAGUUACUGACACAGUAAAAGGAGAACGGGUAUCGUUCCCUGUGUGCACGUACAUC<br>CCGGCCACCAUAUGCGAUCAGAUGACUGGUAUAAUGGCCACGGAUAUAUCACCUGACGAUGCACAAAAACUU<br>CUGGUUGGGCUCAACCAGCGAAUGCUUCUUGCUAUGCAAACUAACAGGAACACCAACACCAUGCAAAAUUAC<br>CUUCUGCCGAUCAUAGCACAAGGGUUCAGCAAAUGGGCUAAGGAGCGCAAGGAUGAUCUUGAUAACGAGAAA<br>AUGCUGGGUACUAGAGAACGCAAGCUUACGUAUGGCUGCUUGUGGGCGUUUCGCACUAAGAAAGUACAUUC<br>GUUUUAUCGCCCACCUGGAACGCAGACCUGCGUAAAAGUCCCAGCCUCUUUUAGCGCUUUUCCCAUGUCGU<br>CCGUAUGGACGACCUCUUUGCCCAUGUCGCUGAGGCAGAAAUUGAAACUGGCAUUGCAACCAAAGAAGGAG<br>GAAAAACUGCGACCUGUCAUUAACGGUAGGAUUAGUCAUGGAGGCCAUUUUCCGAAUCGGAUUUUGUUUUU<br>AAUAU |
